# RD5-mediated lack of PE_PGRS and PPE-MPTR export in BCG vaccine strains results in strong reduction of antigenic repertoire but little impact on protection

**DOI:** 10.1101/265462

**Authors:** Louis S. Ates, Fadel Sayes, Wafa Frigui, Roy Ummels, Merel P. M. Damen, Daria Bottai, Marcel A. Behr, Wilbert Bitter, Laleh Majlessi, Roland Brosch

**Author notes:** Corresponding authors: Louis S. Ates, and Roland Brosch.

## Abstract

Tuberculosis is the deadliest infectious disease worldwide. Although the BCG vaccine is widely used, it does not efficiently protect against pulmonary tuberculosis and an improved tuberculosis vaccine is therefore urgently needed. *Mycobacterium tuberculosis* uses different ESX/Type VII secretion (T7S) systems to transport proteins important for virulence and host immune responses. We recently reported that secretion of T7S substrates belonging to the mycobacteria-specific Pro-Glu (PE) and Pro-Pro-Glu (PPE) proteins of the PGRS (polymorphic GC-rich sequences) and MPTR (major polymorphic tandem repeat) subfamilies required both a functional ESX-5 system and a functional PPE38/71 protein for secretion. Inactivation of *ppe38/71* and the resulting loss of PE_PGRS/PPE-MPTR secretion were linked to increased virulence of *M. tuberculosis* strains. Here, we show that a predicted total of 89 PE_PGRS/PPE-MPTR surface proteins are not exported by certain animal-adapted strains of the *M. tuberculosis* complex including *M. bovis.* This Δ*ppe38/71*-associated secretion defect therefore also occurs in the *M. bovis*-derived tuberculosis vaccine BCG and could be restored by introduction of the *M. tuberculosis* ppe38-locus. Epitope mapping of the PPE-MPTR protein PPE10, further allowed us to monitor T-cell responses in splenocytes from BCG/M. *tuberculosis* immunized mice, confirming the dependence of PPE10-specific immune-induction on ESX-5/PPE38-mediated secretion. Restoration of PE_PGRS/PPE-MPTR secretion in recombinant BCG neither altered global antigenic presentation or activation of innate immune cells, nor protective efficacy in two different mouse vaccination-infection models. This unexpected finding stimulates a reassessment of the immunomodulatory properties of PE_PGRS/PPE-MPTR proteins, some of which are contained in vaccine formulations currently in clinical evaluation.

## Introduction

Tuberculosis is the deadliest infectious disease worldwide and is responsible for more than 1.7 million deaths per year [1]. Its causative agent, *Mycobacterium tuberculosis*, is a slow growing bacterium inherently resistant to many antibiotics. This problem is further exacerbated by rising levels of acquired drug resistance, resulting in multi-drug-resistant (MDR) and extensively-drug-resistant (XDR) strains of *M. tuberculosis,* which require treatment regimens of two years with low treatment success rates and severe side effects [1–3]. These worrying developments highlight the need for a successful vaccine, halting the transmission of tuberculosis [4]. The currently used vaccine is based on of *Mycobacterium bovis*, attenuated through serial culture by Calmette and Guérin and therefore known as Bacille Calmette-Guérin (BCG) [5–7]. BCG is generally believed to protect relatively well against severe forms of disseminated tuberculosis in children, but is unable to induce full protection or halt transmission of *M. tuberculosis* in adolescents and adults [4,8,9]. Furthermore, even these protective traits are subject to controversy, which may be caused by the plethora of genomic mutations and recombination events that have accrued during the worldwide sub-culturing of the original BCG strain [5,6,10,11].

One possible reason for sub-optimal protection by BCG and other candidate vaccines is the absence or secretion defect of certain immunogenic proteins. *M. tuberculosis* secretes many proteins through its different secretion systems, including Sec-translocation (Sec), Twin-arginine-translocation (Tat), or Type VII secretion (T7S) systems [12,13]. *M. tuberculosis* possesses five different T7S systems called ESX-1 to ESX-5 [14]. The first T7S system to be discovered was ESX-1, identified by the Region of Difference (RD)1 deletion in BCG [15], responsible for the loss of ESX-1-mediated secretion in this vaccine strain [16,17]. Substrates of the ESX-1 system are responsible for the rupture of mycobacterium-containing phagosomes and represent a major virulence factor of pathogenic mycobacteria [18–21]. Corresponding to this information, the expression of the ESX-1 secretion system in BCG increased protective activity, but was also associated with increased pathogenesis [22]. Interestingly, a recently developed recombinant BCG strain expressing ESX-1 of *Mycobacterium marinum* was able to induce cytosolic pattern recognition and better protective responses, without a significant increase in virulence [23]. Similarly, the vaccine candidate MTBVAC was recently shown to induce immune responses to selected ESX-1 substrates and this ability was found to be the major determinant of improved protective efficacy as compared to BCG [24].

While the ESX-1 system is the best studied T7S system in mycobacteria, the ESX-5 system has the largest repertoire of substrates [25–27]. The ESX-5 system is essential for slow-growing mycobacteria, because of its role in outer membrane permeability [26,28]. Therefore, this system is present and considered functional in BCG. The coding sequences of the potential substrates of the ESX-5 system together form almost 8% of the coding potential of the *M. tuberculosis* genome [29]. Most notable amongst the ESX-5 substrates are the PE and PPE proteins, named for the proline and glutamic acid residues in their N-terminal domains. Defined functions have been described for some PE-PPE proteins, such as the lipase LipY [30,31] and PPE10, the latter of which is important for capsular integrity of *M. marinum* [32]. Furthermore, many studies have ascribed immunomodulatory functions to PE-PPE proteins, such as altering host cytokine responses by interaction with Toll-like receptors or inhibition of antigenic presentation [33–36]. However, most PE and PPE proteins have no known functions and their high degree of homology makes them difficult to study. The latter is especially true for the two most-recently evolved subgroups of ESX-5 substrates, *i.e.* the PE_PGRS and PPE-MPTR proteins. Both these sub-groups are characterized by their GC-rich DNA sequences, repetitive glycine-rich amino acid motifs and high molecular weight ranging up to ∼4500 kDa [27,29].

We recently identified the PPE protein PPE38 and its highly similar duplicated variant PPE71, as essential factors in the secretion of both the PE_PGRS and PPE-MPTR proteins, in both *M. marinum* and *M. tuberculosis* [37]. The genes encoding PPE38 and PPE71 are organized in a 4-gene locus that also includes the *esxX* and *esxY* genes (Figure1A), which however are not required for PE_PGRS secretion in *M. tuberculosis* strain CDC1551 [37]. Strains with naturally occurring, or engineered, loss-of-function mutations of the ppe38-locus were unable to secrete both PE_PGRS and PPE-MPTR proteins and were more virulent in a mouse infection model [37]. Indeed, deletion of the ppe38-locus occurred at the branching point of modern Beijing (Lineage 2) strains and may have aided in their global dispersal [37]. Moreover, the ppe38-locus was previously shown to be a hypervariable genetic region and many strains within the *M. tuberculosis* complex (MTBC) have polymorphisms in this locus [38]. The most well-known of these polymorphisms is the deletion of the RD5 region from BCG and several other animal-adapted strains of the MTBC [38,39].

**Figure 1:**
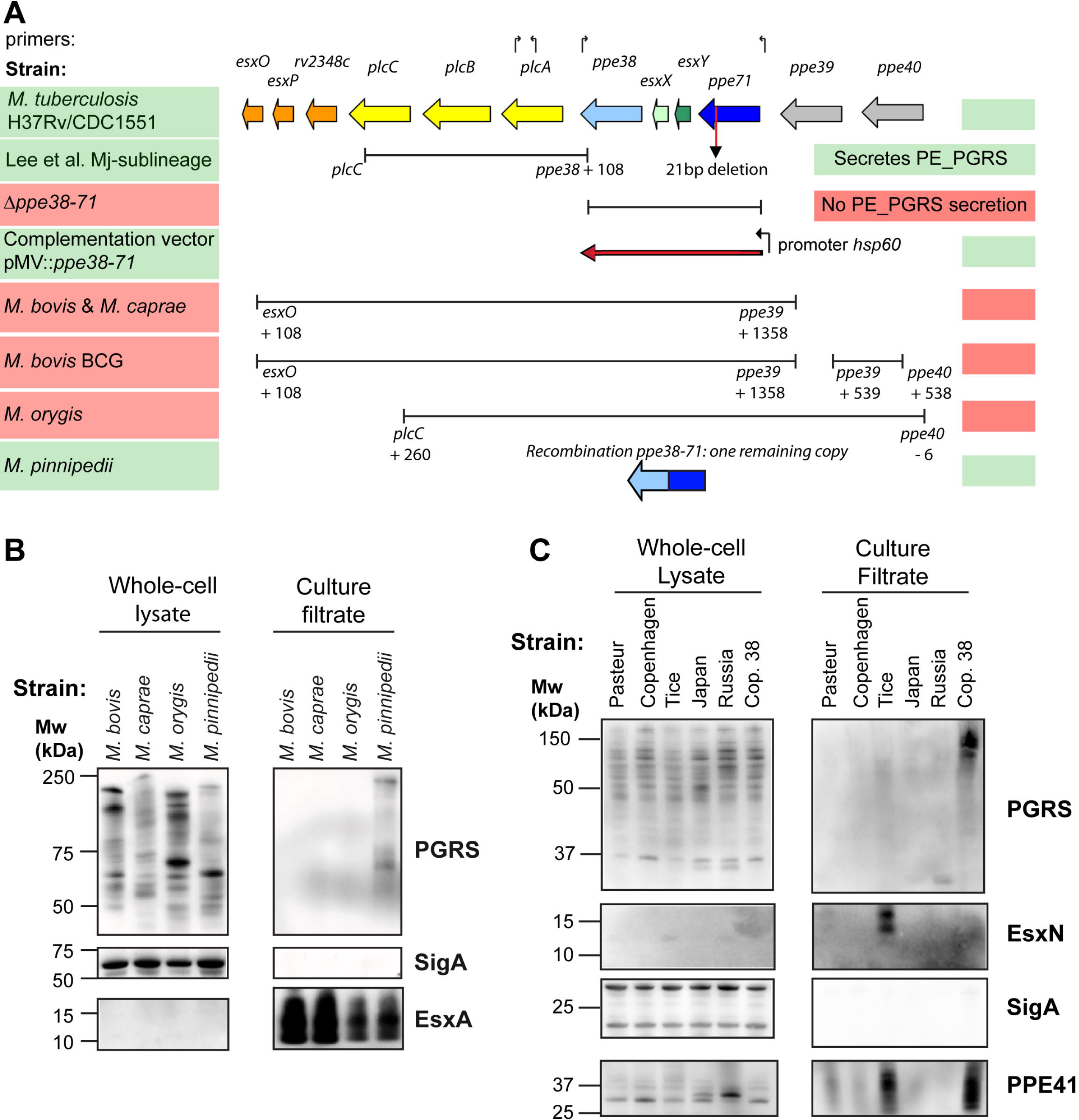
RD5-like genetic deletions in the *M. tuberculosis* complex and their effect on PE_PGRS secretion. A) The genetic organization of the RD5 locus in *M. tuberculosis* strains CDC1551 and H37Rv is depicted in colored arrows. Bars below the genes indicate the size and location of different RD5-like and *ppe38*-deletions examined in this work. Arrows above the genes indicate primers used in this study to verify the presence of RD5 associated genes, sequences can be found in Supplementary Table 4. Functional PE_PGRS secretion is indicated by shading of the strain name in green, while red shading represents strains in which PE_PGRS secretion is not functional (based on immunoblot analysis). Figure adapted from Mc Evoy et al. 2009 with permission from the authors [38]. B) Immunoblot secretion analysis of animal-adapted MTBC strains verifies that strains with RD5 deletions do not secrete PE_PGRS proteins. C) Immunoblot secretion analysis of five genetically divergent BCG isolates confirms the PE_PGRS secretion defect in all BCG isolates. Cop. 38 indicates the strain *M. bovis* BCG Copenhagen, transformed with vector pMV::*ppe38-71* and is hereafter referred to as BCG38. Full western blots corresponding to panels depicted in B-C are depicted in Supplemental Figure 5.

The biological impact of the RD5 deletion has been a controversial subject of research and has focused solely on the phospholipase C encoding genes *plcABC.* Deletion of *plcABC* was reported to either attenuate [40] or increase virulence of *M. tuberculosis* [41]. However, a more recent study of the plc-genes in different mouse and cellular models showed no relevant contribution of these genes to the virulence of *M. tuberculosis* [42].

Here, we investigated the effect of RD5-like polymorphisms of the *ppe38-locus* in a number of MTBC-branches and discovered that the RD5 deletions in animal-adapted strains and the BCG vaccine strains have profound effects on the repertoire of secreted substrates in these strains. Restoration of PPE38-dependent secretion results in a wider antigenic repertoire of BCG, whereby the identification of two immunogenic epitopes in one of the substrates, *i.e.* the PPE-MPTR protein PPE10, has allowed us to monitor the immunological impact of the corresponding secretion characteristics on host immune responses.

## Results

### Variation in PE_PGRS secretion in MTBC lineages and outgroups indicate genome sequence errors

The genetically most-distant tubercle bacilli are represented by the *Mycobacterium canettii* clade. This outgroup mirrors the genomic diversity likely present within the ancestor of *M. tuberculosis* before branching and clonal expansion of the MTBC [43]. Recent studies of *M. canettii* have improved our understanding of adaptations that have shaped the transition from an *M. canettii-like* ancestor into extant *M. tuberculosis*, such as the gain of surface hydrophobicity through loss of lipooligosaccharide production [44] and the apparent loss of the capacity to exchange chromosomal DNA in the MTBC [45]. Interestingly, the available genome sequence information of five *M. canettii* isolates revealed potential polymorphisms in the *ppe38-locus* [43]. While strains D, K and L all possessed copies of the *ppe38* and *ppe71* genes, the sequence of strain J in the database indicated the potential absence of *ppe38* and *ppe71* from the strain. Such a deletion would be expected to affect PE_PGRS secretion [37]. However, secretion analysis revealed that all 5 isolates secreted PE_PGRS proteins (Supplemental Figure 1A). Subsequent PCR analysis confirmed the presence of a complete *ppe38-71* locus, similar to *M. tuberculosis* H37Rv, for all tested *M. canettii* strains, including strain J (Supplemental Figure 1B). It is likely that the sequence polymorphisms in the previously deposited dataset may have arisen due to automated sequence assembly-associated bio-informatic artefact, which is a known problem for this region [37,38].

Another interesting group of strains, which were reported to have major polymorphisms in the RD5/ppe38-locus, was recently described by Lee et al. [46]. The Inuit population of the Nunavik region in Canada is affected by high levels of tuberculosis incidence. The majority of all cases in this cohort were shown to have resulted from the introduction of a single, particular *M. tuberculosis* strain, about one century ago. This sublineage was defined by genomic deletions, two of which affect the *ppe38* locus. A 5,759bp RD5-like deletion removed the three phospholipase C genes *plcABC* and truncated *ppe38* (Figure 1A). The other *ppe* gene in this locus, *ppe71* (*mt2422*), was reported to be affected by 22bp frameshift deletion (Figure 1A)[46]. Reinvestigation of the sequence of *ppe71* by inspection of the whole genome sequence data, and by PCR and Sanger sequence analysis, revealed that this deletion was in fact a 21bp deletion causing a 7 amino-acid deletion (PPE38 Amino acid 354-MGGAGAG-361) instead of a frameshift. This deletion has been previously described to occur also in other strains of *M. tuberculosis*, including CDC1551 (MT2422 - http://www.genome.jp/dbget-bin/www_bget?mtc:MT2422) [38]. To test whether the RD5-like polymorphism (CDC1551-D17) negatively affects PE_PGRS secretion, five strains with and one strain without this deletion were subjected to secretion analysis by immunoblotting. All strains exhibited similar secretion levels of both PE_PGRS proteins and the ESX-1 substrate EsxA, as compared to reference strain CDC1551 (Supplemental Figure 1B). These data show that the PPE71 variant carrying the MGGAGAG-deletion is able to sustain PE_PGRS secretion levels in *M. tuberculosis*, independently of truncation of PPE38. Furthermore, there is no apparent phenotypic difference when *M. tuberculosis* has one or two functional copies of PPE38/71.

### RD5 deletions in animal-adapted strains and in *M. bovis* BCG block PE_PGRS secretion

A striking amount of different RD5-like polymorphisms are present in the animal-adapted lineages/ecotypes of *M. tuberculosis* complex. These strains share their most recent common ancestor with *M. africanum* Lineage 6 [47,48], which is reported to have two copies of *ppe38/ppe71* [49]. *Mycobacterium pinnipedi*, a pathogen for seals and sea lions, has one intact copy of the *ppe38* gene, but no esxXY-genes (Figure 1A). *M. bovis* and *Mycobacterium caprae* share an identical RD5 deletion, while *Mycobacterium orygis* possesses a unique RD5 deletion (Figure 1A) [38,50,51]. To investigate the effect of RD5 deletions on PE_PGRS secretion in animal-adapted strains, we performed secretion analysis of *M. bovis, M. caprae, M. orygis* and *M. pinnipedi* (Figure 1B). As expected, *M. pinnipedi* was the only tested species able to secrete PE_PGRS proteins in concordance with the presence of one functional copy of *ppe38* (Figure 1B). In contrast, *M. bovis, M. caprae* and *M. orygis* were deficient in PE_PGRS secretion, while EsxA secretion was not affected and no marked cell lysis occurred (Figure 1B). Intracellular PE_PGRS expression was detected in strains with a secretion defect and was strikingly different between isolates (Figure 1B).

Since *M. bovis* and *M. caprae* share the same RD5 deletion with *M. bovis* BCG, we hypothesized that this vaccine strain is also deficient in PE_PGRS secretion (Figure 1A). Indeed, five different *M. bovis* BCG isolates, which were selected for their relative genetic distance [6,10], were all deficient in PE_PGRS secretion (Figure 1C). It is of interest to note that *M. bovis* BCG Tice secretes higher levels of the ESX-5 substrates PPE41 and EsxN (Figure 1 C), likely because of its genetic duplication of the *esx-5* genetic locus [10]. However, despite this ESX-5 duplication, BCG Tice is unable to secrete PE_PGRS proteins. The PE_PGRS secretion defect of BCG was not restored in a previously constructed BCG strain with a cosmid containing the complete RD5 region of *M. tuberculosis* H37Rv (Supplemental Figure 1C) [16]. In contrast, introduction of the *ppe38-71* locus from *M. tuberculosis* on an integrative plasmid constitutively expressing these genes under control of the *hsp60* promoter [37], partially restored PE_PGRS secretion of recombinant *M. bovis* BCG. This finding was especially surprising since emergence of RD5-deleted *M. bovis/M. caprae* progenitor strains likely dates back thousands of years [47]. The obtained *ppe38-71*- complemented *M. bovis* BCG Copenhagen strain was named BCG38.

Taken together, our data show that the different BCG vaccines are all deficient for the secretion of PE_PGRS proteins and that this is at least partly revertible by complementation with the *ppe38-71* locus of *M. tuberculosis.* Based on our previous work, this secretion defect is expected to affect up to 89 proteins classified as PE_PGRS or PPE-MPTR [27,37].

### Secretion of PE_PGRS/PPE-MPTR proteins in *M. tuberculosis* or BCG does not alter phenotypic and functional maturation of host innate immune cells, or antigenic presentation

The ability to restore PPE38-dependent secretion in *M. bovis* BCG allowed us to investigate to what extent this secretion defect affects properties of the BCG vaccine. Many of the 89 members of the PE_PGRS and PPE-MPTR proteins have been suggested to perform biological roles in virulence and immune modulation, although the molecular mechanisms and biological relevance remain unestablished for most of these [14,27,33,34]. Increasing the repertoire of immunogenic proteins secreted by BCG could lead to increased protection, since protein secretion by mycobacteria is essential for the efficient induction of protective CD4+ T-cell responses [22,52–55]. However, restoring secretion of proteins that have been proposed to exhibit immunomodulatory functions could also decrease efficacy of the vaccine strain. In particular, recent reports suggest that PPE38 itself downregulates Major Histocompatibility Complex class-I (MHC-I) expression in murine macrophages [56] and that PE_PGRS47 inhibits autophagy and is responsible for reducing MHC-II-restricted antigen presentation during *in vivo* infection of mice [35].

We set out to establish whether presence of PPE38 and the ability to secrete PE_PGRS and PPE-MPTR proteins, affected phenotypic and functional maturation of infected murine innate immune cells. Bone marrow-derived dendritic cells (BM-DCs) of C57BL/6 mice were infected (MOI = 0.5) with i) *M. tuberculosis* CDC1551, ii) an isogenic deletion-mutant of the complete *ppe38-71* locus (Δ*ppe38-71*), iii) or a complemented strain (*ppe38-71-C*) [37] and were tested in parallel with *M. bovis* BCG Copenhagen and the isogenic strain expressing *ppe38-71* (BCG38). All the infected BM-DCs exhibited a clear upregulation of co-stimulatory markers CD40, CD80 and CD86, as well as modulation of MHC-I (H-2K^b^) and MHC-II (I-A^b^) expression, compared to uninfected controls. However, no differences in the induction of any such phenotypic maturation markers could be observed for the different isogenic WT and recombinant strains (Figure 2, Supplemental Table 1). Quantification of several inflammatory cytokines in the culture supernatants of the infected BM-DCs showed highly similar levels of TNFα, IL-12p40/70 and IL-6 production induced by the isogenic strains of BCG and *M. tuberculosis* (Figure 2B). These results indicate that PPE38-dependent secretion defects are unlikely to have a major effect on the phenotypic or functional maturation of DCs, even though many PE_PGRS and PPE-MPTR proteins have previously been suggested to perform such biological roles [33,35].

**Figure 2:**
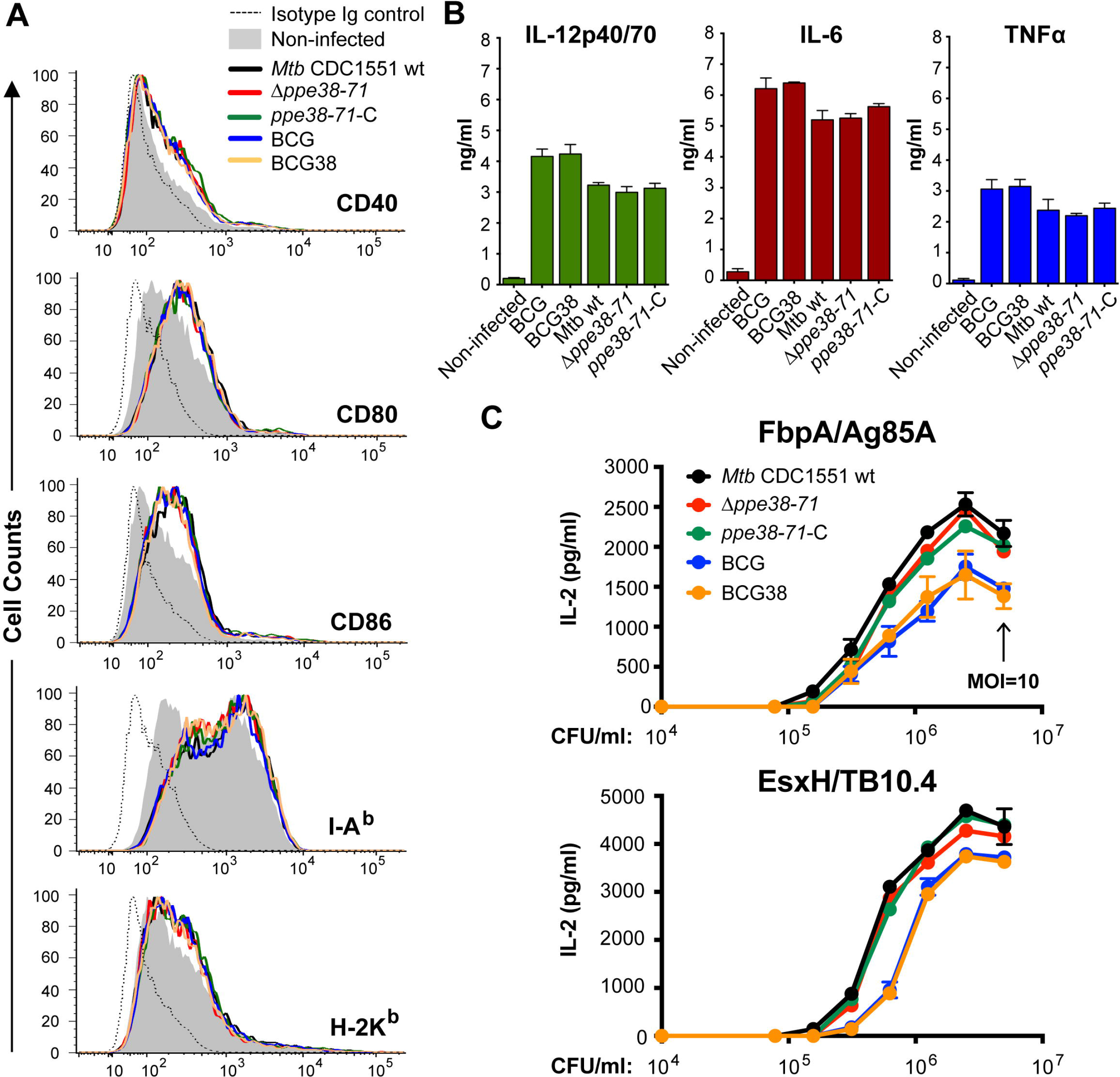
Secretion of PPE38 and PE_PGRS/PPE-MPTR proteins in BCG or *M. tuberculosis* does not alter phenotypic and functional maturation, or antigen presentation by innate immune cells. A) BM-DCs (C57BL/6, H-2^b^) infected with the indicated mycobacterial strains were stained for surface expression of co-stimulation markers CD40, CD80 and CD86, or MHC components I-A^b^ and H-2K^b^. Depicted are the cell counts (Y-axis) and fluorescent intensity (X-axis) as quantified by flow cytometric analyses. Quantification of mean fluorescent intensity and quantification of cell survival can be found in Supplemental Table 1. B) Culture supernatant of the experiment described in A was assessed for the presence of cytokines IL-12p40/70, IL-6 and TNF-α. No differences were detected between cells infected with the isogenic BCG or *M. tuberculosis* isolates. C) Antigenic presentation by infected DCs is not affected by disruption or restoration of PPE38–dependent protein secretion in *M. tuberculosis* or BCG. BM-DCs (BALB/c, H-2^d^) were infected with two-fold dilutions (data points in graph) of the indicated *M. tuberculosis* or BCG strains starting at MOI=10 (indicated by black arrow). IL-2 production was quantified by ELISA after overnight co-culture with I-E^d^-restricted T-cell hybridoma specific for FbpA (Ag85A_101-120_ (2A1), upper panel) or with I-A^d^-restricted T-cell hybridoma specific for EsxH (TB10.4_74-88_ (1G1), lower panel). Data are representative of biological duplicates.

In addition, we assessed whether PPE38-dependent protein secretion influences MHC-II-restricted presentation of other mycobacterial antigens. Such a phenotype might possibly be caused by a direct effect on the host phagocytes due to restored PE_PGRS secretion [35,36], or by competition in the hosts antigen presentation machinery upon secretion of the large number of PPE38-dependent substrates. To test this hypothesis, BM-DCs were infected with serial two-fold dilutions of *M. tuberculosis* (CDC1551) or the isogenic Δ*ppe38-71* deletion or complemented strains, as well as BCG or BCG38. MHC-II restricted T-cell hybridomas specific to FbpA (Ag85A_101-120_) or EsxH (TB10.4_74-88_) T-cell epitopes were added after overnight infection and washing. IL-2 secretion in culture medium was quantified by ELISA as a measure of antigen presentation and hybridoma T-cell activation. T-cell hybridomas specific for both FbpA (Figure 2C, upper panel) and EsxH (lower panel) produced higher levels of IL-2 in response to *M. tuberculosis* strains compared to BCG strains. However, no differences were observed between isogenic strains with, or without, functional PPE38-dependent PE_PGRS/PPE-MPTR secretion. These data show that PPE38-dependent PE_PGRS/PPE-MPTR secretion does not reduce MHC-II-restricted antigen presentation of other mycobacterial antigens by the host DCs.

Together, these results suggest that introduction of PPE38 and restoration of PE_PGRS secretion do not negatively affect phenotypic and functional maturation of innate immune cells, or their capacity to present antigen to CD4^+^ T cells.

### Restoration of PPE38-dependent PE_PGRS/PPE-MPTR protein secretion in BCG does not impact protection potential against *M. tuberculosis* in mice

Since we found no evidence suggesting that antigen presentation of mycobacterial antigens by DCs is negatively affected by restoration of PPE38-dependent secretion, we hypothesized that the enlarged repertoire of secreted proteins in BCG38 could increase its vaccine potential compared to the parental BCG. In parallel, we hypothesized that the capsule of BCG could be altered upon restoration of PPE38-dependent secretion. We recently reported that transposon insertions in the gene encoding an ESX-5 associated chaperone (*espG_5_*), or in the PPE-MPTR encoding gene *ppe10* (*mmar_0761*), reduce capsule integrity of *M. marinum* [32]. Similarly, an *eccC_5_::tn* mutant in the *M. tuberculosis* strain CDC1551, completely deficient in ESX-5 secretion, also exhibited reduced capsule integrity [32,57]. Since PPE10 is dependent on PPE38 for its secretion [37], we hypothesized that restoration of PPE10 secretion might positively affect capsule integrity. The presence of an intact capsule on BCG, achieved by culturing in detergent-free growth medium, has recently been shown to be important for a more potent immune response and could therefore be relevant for the protective efficacy of BCG38 [58].

To test both hypotheses, C57BL/6 mice were subcutaneously (s.c.) immunized with 1 million CFU of either BCG, or BCG38, cultured either in shaking condition in the presence of 0.025% Tween-80, or in unperturbed conditions without detergent. Four weeks post-immunization, mice were challenged by an aerosol infection of *M. tuberculosis* H37Rv (bacterial load: 680 CFU/lung at Day 1, prepared without detergent). Mice were killed four weeks post infection, at which time lungs and spleens were harvested and assessed for bacterial burdens by CFU counting. An approximate 100-fold reduction in bacterial lung burdens was achieved by all conditions of vaccination irrespective of the presence of detergent, or the BCG *vs* BCG38 vaccine strains (Figure 3A). This reduction of bacterial lung burden coincided with improved macroscopic state of the lungs (Supplemental Figure 2A). Similarly, an approximately 10-fold reduction in spleen CFUs and reduction in splenomegaly was detected in the vaccinated mice irrespective of the method of vaccine preparation (Figure 3A, Supplemental Figure 2B). No significant (*p*<0.05) differences in bacterial burdens were observed between any of the four tested conditions in either the spleens or lungs. Together, these results show that restoration of PPE38-dependent PE_PGRS/PPE-MPTR secretion in BCG does not significantly improve protection against *M. tuberculosis* in the murine model used. Moreover, we did not find a significant difference in protective efficacy between conventional and detergent-free preparation of either BCG or BCG38, suggesting that capsular integrity is not altered or does not affect protection in this model.

**Figure 3:**
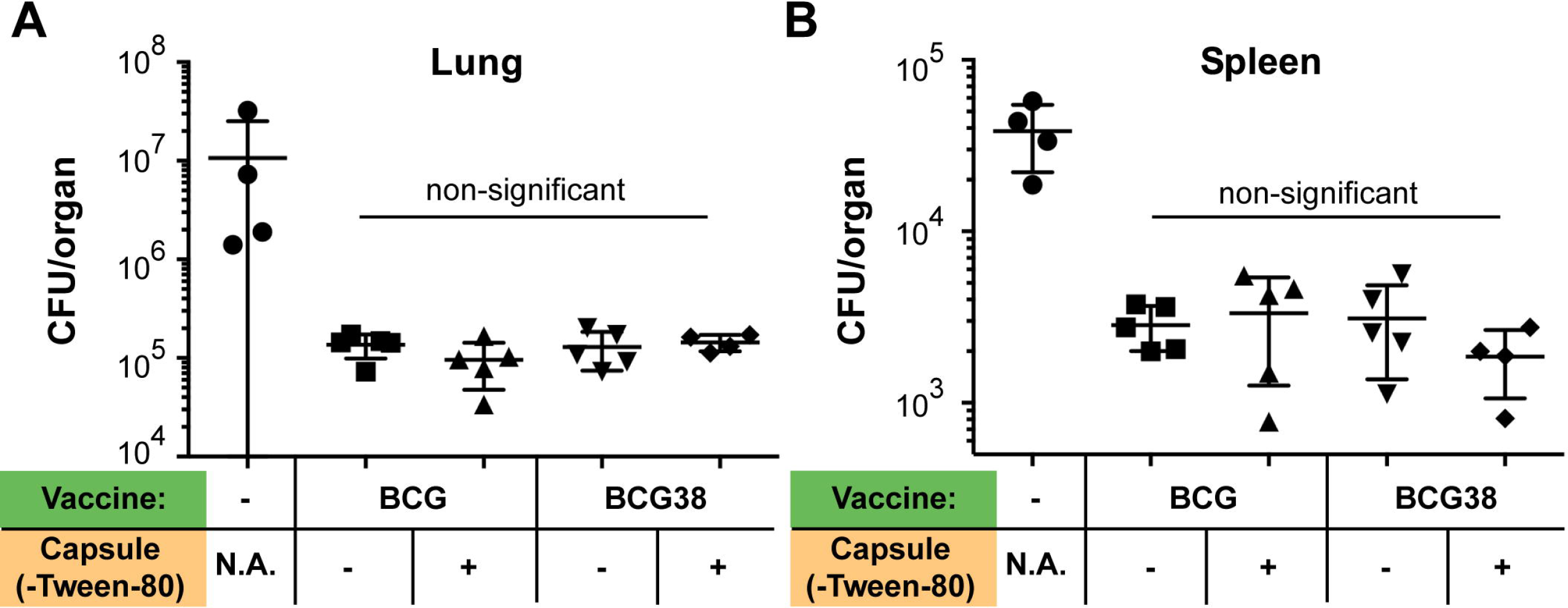
Restoring PPE38-dependent protein secretion of BCG does not increase protection against *M. tuberculosis* in C57BL/6 mice. Lung (A) or spleen (B) bacterial burdens of C57BL/6 mice infected with *M. tuberculosis* H37Rv via aerosol administration. Mice were vaccinated s.c. four weeks before the challenge, with 1 × 10^6^ CFU/mouse of either BCG or BCG38 (indicated in green). Both strains were prepared, either in standard culture conditions in medium containing 0.025% Tween-80 considered as no capsule (indicated with (−), or in culture allowing capsule formation/retention in detergent free condition (indicated with (+)). Photographs of the assessed organs are depicted in Supplemental Figures 2A, B. Each data point represents the CFU/organ of one single mouse counted and averaged from two technical duplicates. Error bars depict the standard deviation. Differences between different vaccination conditions were non-significant (*p*>0.05), but all vaccination conditions were statistically different from the unimmunized control group (*p*<0.01). Significance was calculated with Prism software using ordinary one-way ANOVA followed by Tukey’s test for multiple comparisons.

### Identification of immunogenic T-cell epitopes of the PPE-MPTR protein PPE10

Secretion of T7S-mediated mycobacterial proteins is essential to induce host CD4+ T-cell responses and the great majority of immunogenic and protective antigens of *M. tuberculosis* are secreted proteins [59]. Many of the known immunodominant antigens are PE and PPE proteins and these form an integral part in a number of subunit or recombinant vaccines [52,60–63]. Therefore, the finding that restoration of PPE38-dependent PE_PGRS/PPE-MPTR secretion in BCG did not significantly affect protective efficacy was surprising, particularly as up to 89 individual proteins are predicted to be concerned. In order to explain these unexpected data, we reflected on our hypotheses and found additional variables that could affect the assumptions on which they are based. In particular, while PPE-MPTR secretion was shown to be strictly dependent on PPE38 in both *M. marinum* and *M. tuberculosis*, we had no direct evidence of PPE-MPTR secretion in BCG38. In contrast to PPE-MPTR proteins, PE_PGRS proteins may not contain immunodominant epitopes or be protective antigens [64–67]. Furthermore, although previous studies have found a strict correlation between *in vitro* secretion and the capability to induce CD4+ T-cell responses [23,52,63], it is conceivable that the PPE38-dependant substrates are still membrane, or surface, associated in *ppe38-71*-deficient strains and thereby remain able to induce T-cell responses.

Since tools to study PPE-MPTR proteins are scarce and currently insufficient to answer the questions above, we set out to develop an immunological approach to study PPE-MPTR secretion and their immunogenicity in more detail. We selected PPE10 as a model MPTR-protein, because PPE10 is predicted to be the most ancestral MPTR protein in mycobacteria [27]. The PPE domain covers the N-terminal 181 residues of PPE10 and is highly similar to other PPE proteins. The middle of the protein contains a typical MPTR repeat domain, which is very similar to other MPTR proteins. The C-terminus contains a domain unique to PPE10, which is secreted *in vitro* [25,32,37]. PPE10 is also of biological interest, since it is detected *in vivo* in guinea pig lungs and this protein is required for capsular integrity of *M. marinum* [32,68]. We set out to assess whether PPE10 has the potential to induce CD4+ T-cell mediated immune responses in mice. To increase the likelihood of identifying immunogenic epitopes, we immunized not only C57BL/6 mice, but also C57BL/6 x CBA (H-2^b/k^) F1 mice, which express a more diverse repertoire of MHC restricting elements. While C57BL/6 mice only express a single MHC-II molecule (I-A^b^), C57BL/6 x CBA F1 mice can potentially express six different MHC-II variants (Supplemental Table 2). Mice were s.c. immunized with wild-type *M. tuberculosis* H37Rv and were killed three weeks later. Splenocytes were isolated and stimulated *in vitro* with a peptide library consisting of sixty 15-mers with a 5-amino acid shifting frame spanning PPE10_181-487_ of *M. tuberculosis* H37Rv [29,69]. None of the sixty peptides were able to induce specific T-cell mediated IFN-γ responses by splenocytes from unimmunized mice or immunized C57BL/6 mice (Supplemental Figure 3). However, two peptides were immunogenic in the C57BL/6 x CBA (H-2^b/k^) F1 mice and induced high levels of IFN-γ, similar to the positive control peptide ESAT-6_1-20_ (Figure 4, Supplemental Figure 3). Interestingly, one of these immunogenic peptides (PPE10_221-235_: GSGNTGSGNLGLGNL) was situated in the MPTR domain of PPE10, while the other (PPE10_381-395_: NVLNSGLTNTPVAAP) was derived from the PPE10-specific C-terminal domain. The MPTR peptide PPE10_221-235_ has 17 close homologues within the *M. tuberculosis* genome (identity > 65%, but no 100% homologues), while this was not the case for PPE10_381-395_ (Supplemental Table 3). These results show that immunization with *M. tuberculosis* induces immune responses against PPE10 and that this response can be elicited both against the PPE10-specific C-terminal domain or the MPTR domain.

**Figure 4:**
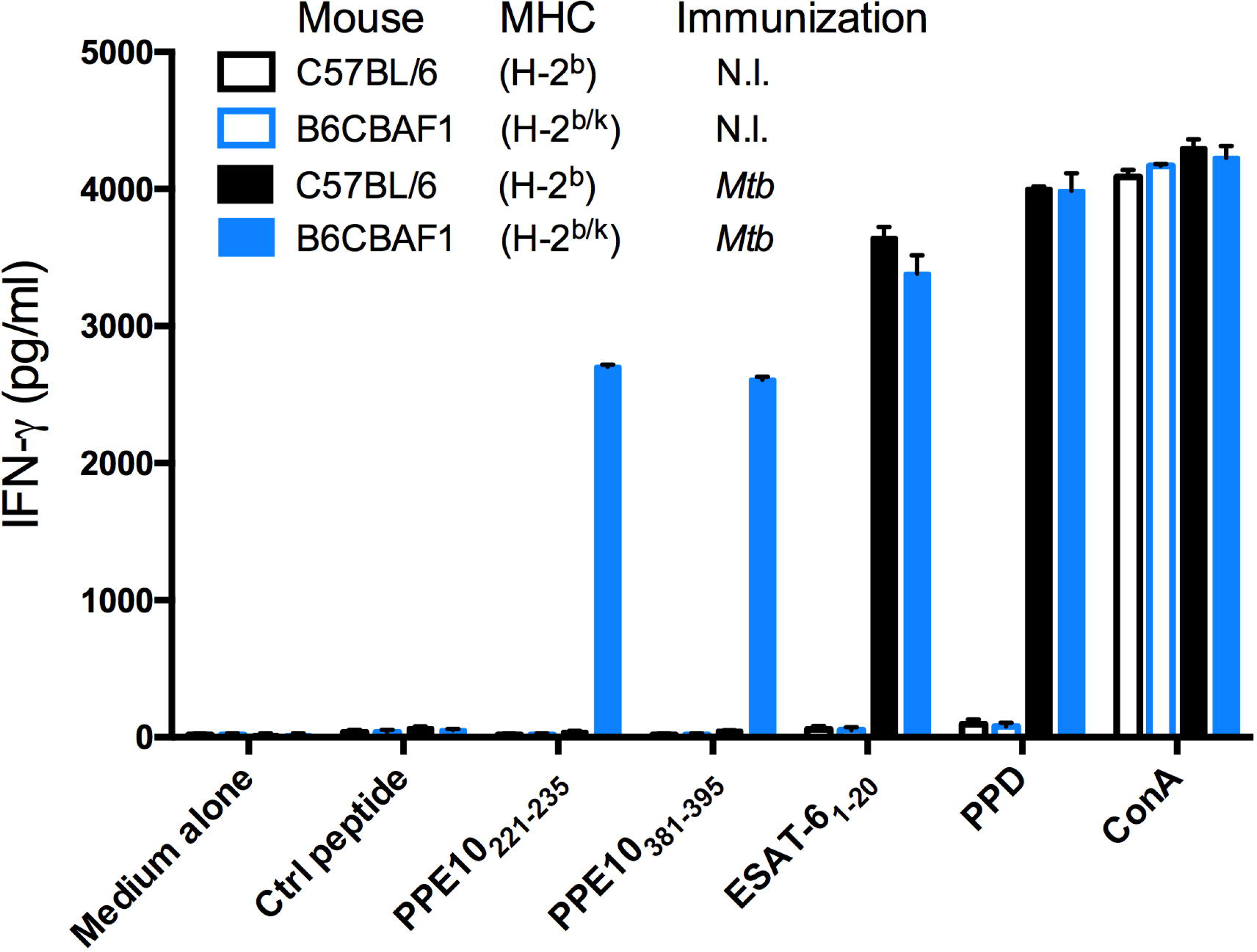
Epitope mapping of PPE10 identifies two novel immunogenic T-cell epitopes. C57BL/6 H-2^b^ (black) or C57BL/6 x CBA (H-2^b/k^) F1 mice (B6CBAF1, blue) were immunized s.c. with 1 × 10^6^ CFU/mouse of *M. tuberculosis* H37Rv (*Mtb*, filled bars), or were left non-immunized (N.I. empty bars). Three weeks post-immunization, splenocytes were restimulated with control peptides or a library of 15-mers spanning PPE10 excluding the PPE domain. T-cell mediated IFN-γ responses were quantified buy ELISA as a measure of immunogenicity. Two immunogenic PPE10-peptides were identified (PPE10_221-235_ & PPE10_381-395_) in B6CBAF1 mice. Error bars depict standard deviation over two technical replicates. This figure depicts only newly identified epitopes and controls. Full results of the pep-scan epitope mapping can be found in Supplemental Figure 2.

### Deletion of *ppe10* does not significantly alter protein secretion of other Type VII secretion substrates in *M. tuberculosis*

The newly identified immunogenic peptides derived from PPE10 are a tool that allowed us to answer different questions regarding the PPE-MPTR proteins. First, to determine the specificity and cross-reactivity of the epitopes, we constructed a deletion mutant of *ppe10* (*Rv0442c*) in the *M. tuberculosis* CDC1551 background by homologous recombination and phage transduction (Supplemental Figure 4) [70]. In contrast to *M. marinum-ppe10::tn* [32], no altered colony morphology or other growth phenotype was observed in *M. tuberculosis*-Δ*ppe10*. This finding is in concordance with the absence of such a phenotype in ESX-5 mutants of *M. tuberculosis* and highlights this as a species-specific difference between *M. marinum* and *M. tuberculosis* [32,57,71].

We performed biochemical secretion analysis on the Δ*ppe10* strain in parallel with the strains that were examined for their immunogenic potential (see below). This secretion analysis confirmed the expected PE_PGRS secretion defects of BCG, *M. tuberculosis*-Δ*ppe38-71* and *eccC5::tn*, which were restored in the complemented strains, *i.e. ppe38-71-C* and BCG38 (Supplemental Figure 4C) [37,57]. As expected, BCG and BCG38 were deficient in secretion of the ESX-1 substrate EsxA (ESAT-6) and exhibited only low levels of PPE41 and EsxN. The increase of PPE41 secretion in BCG38 compared to the parental strains (Figure 1C) was consistent in this experiment and other replicates. In contrast to a previous report, we found that *M. tuberculosis* Δ*ppe25-pe19* did secrete PPE41 and EsxN, which may be due to differences in bacterial growth conditions and/or methods in protein extraction and detection [71]. This strain harbors intact genes coding for the ESX-5-membrane complex [57,72] and is able to induce *in vivo* CD4*+* T-cell responses against PE and PPE proteins, in contrast to the general ESX-5 deficient strain Δ*eccD_5_* in the same background [52,63]. The Δ*ppe10* strain showed no difference in PE_PGRS secretion. Similarly, secretion of EsxA and EsxN was not affected by deletion of *ppe10*. Although slightly elevated levels of PPE41 secretion were observed, we concluded from these combined data that *M. tuberculosis*-Δ*ppe10* does not have a general supersecretion phenotype as was previously reported for *M. marinum-ppe10::tn* [32].

### BCG and *M. tuberculosis*-Δ*ppe38-71* are unable to induce immune responses against PPE10

To assess the specificity of the newly identified PPE10 epitopes and to better understand the effect of the *ppe38*-dependent secretion on immunogenicity, we immunized C57BL/6 x CBA F1 mice with the different *M. tuberculosis* and BCG strains for which the secretion phenotype was characterized (Supplemental Figure 4C). Three weeks post-immunization, splenocytes were collected and stimulated with the PPE10_221-235_ and PPE10_381-395_ peptides, as well as purified protein derivate (PPD - a positive control for immunization by Mycobacteria) and a number of known antigenic peptides derived from proteins secreted via ESX-1 (EsxA_1-20_ [73] and CFP-10_11-25_ [74]), ESX-5 (PE19_1-18_ and PPE25_1-20_ [52]) or the twin-arginine-translocation (TAT) pathway (Ag85A_241-260_) [75,76]. As expected, splenocytes of mice immunized with *M. tuberculosis* CDC1551 produced high levels of IFN-γ after stimulation with PPE10_221-235_, PPE10381-395 or all other immunogenic peptides, but not when incubated with a negative control peptide (*E. coli* MalE_100-114_), or the medium control (Figure 5). Splenocytes from unimmunized mice did not react against any of the peptides or PPD and only produced IFN-γ upon simulation by Concanavalin A (ConA). The Δ*ppe10* deletion strain did not induce IFN-γ production in response to either PPE10_221-235_, or PPE10_381-395_, whereas responses against the other peptides were unaffected (Figure 5). Unexpectedly, this result shows that both of the newly identified PPE10 peptides are highly specific, even though we hypothesized cross-reactivity to occur for PPE10_221-235_, because of the high similarity to other MPTR domains (Supplemental Table 3). As expected, the ESX-5 secretion mutant *eccC_5_::tn* did not induce T-cell responses against the ESX-5 substrates PE19, PPE25 and PPE10, further confirming that the export of these antigens by the ESX-5 secretion system is indispensable for the induction of T-cell immune responses [52,63]. Importantly, Δ*ppe38-71* was not able to induce immunogenicity against either of the PPE10 epitopes, a phenotype that was fully reverted in the complemented strain *ppe38-71-C*. This confirms that secretion and *in vivo* immunogenicity of PPE10 as a model PPE-MPTR protein are dependent on PPE38 in the *M. tuberculosis* CDC1551 background, which we were previously unable to assess. Similar to Δ*ppe38-71*, also BCG was completely unable to induce immune responses against either of the PPE10 epitopes. In contrast, BCG38 induced immunogenicity against both PPE10 epitopes at similar levels to the *M. tuberculosis* isolates. Together, these results clearly confirm that the secretion and *in vivo* immunogenicity of the ancestral PPE-MPTR protein PPE10 is strictly dependent on PPE38. These data also provide evidence that the *in vitro* observed PPE38-dependence of PE_PGRS and PPE-MPTR proteins is a phenotype that can be directly translated to the *in vivo* situation. Here, we show that the vaccine strain BCG is unable to induce T-cell responses against the ancestral PPE-MPTR protein PPE10, because of the deletion of its *ppe38-71*-locus as part of RD5.

**Figure 5:**
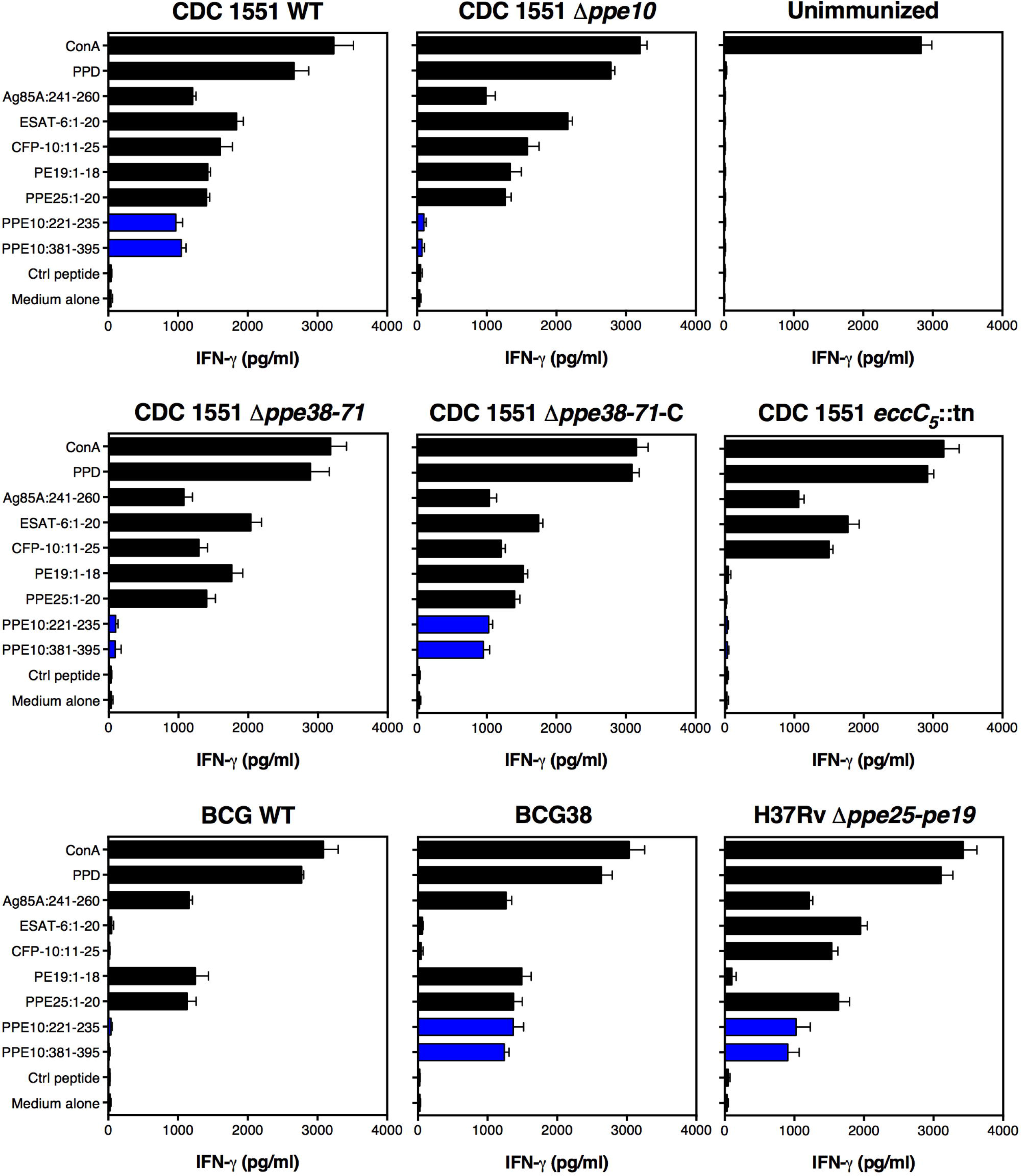
Ability of mycobacteria to induce T-cell responses against PPE-MPTR protein PPE10 is dependent on functional ESX-5- and PPE38-dependent secretion. C57BL/6 x CBA F1 mice were immunized with the indicated mycobacterial strains. Three weeks postimmunization, splenocytes were stimulated with the indicated peptides and IFN-γ production was measured by ELISA. Responses to the newly identified PPE10-derived immunogenic peptides are depicted in blue. Error bars represent the standard deviation over two technical duplicates. The results are representative two biological replicates performed on different timepoints.

Finally, we compared the results obtained for the different WT and recombinant BCG strains with a recently developed attenuated *M. tuberculosis* strain, deleted for 5 *pe/ppe* genes in the *esx-5* locus, named *Mtb*Δ*ppe25-pe19* [71]. Genes encoding the ESX-5 secretion core machinery [57,72] are intact in this strain, as is the *ppe38* gene, a finding which is confirmed by the fact that this strain induced T-cell responses against both PPE10 epitopes. This result highlights that attenuated *M. tuberculosis* vaccine strains may avoid certain *M. bovis* related secretion differences that result in immunogenic properties.

### Prime-boost vaccination regimen to improve PPE10-specific immune responses does not increase protection against *M. tuberculosis*

The results of our epitope mapping analysis showed that C57BL/6 mice were unable to develop T-cell responses against PPE10, which could provide an explanation for the lack of improved protection conferred by BCG38 compared to BCG. Therefore, we set out to perform a similar experiment in these C57BL/6 x CBA F1 mice. In order to maximize any potential increase in PPE-MPTR-specific immune responses, this experiment was designed to boost vaccination of BCG or BCG38 with the immunogenic peptides PPE10_221-235_ and PPE10_381-395_ (Figure 6A). Sixty days after s.c. vaccination with BCG strains, a booster of peptides formulated in the adjuvant CpG(DOTAP), or the adjuvant alone, was administered s.c.. Twenty-nine days later, a second booster was administered intranasally. Nine days after this intranasal boost, mice were challenged by an aerosol challenge of *M. tuberculosis* H37Rv. Mice were killed 28 days later and both lung and spleen bacterial burdens were assessed (Figure 6B, C). No significant differences were observed among the groups of vaccinated animals. Only a modest decrease in spleen CFUs was achieved by any of the vaccination regimens. This reduction was not significant (*p*<0.05) for BCG-vaccinated mice and injected with the adjuvant alone, but was significant for the three other groups (Ordinary one way ANOVA – Dunnett’s test of multiple comparisons against a single control). However, no significant differences between any of the vaccinated groups was detected (Ordinary one way ANOVA, Tukey’s test of multiple comparisons). Vaccination with all regimens reduced lung CFU values at least 10-fold. In fact, vaccination with BCG38, boosted with PPE10-derived immunogenic peptides, had the highest average bacterial lung burden of the four different vaccination regimens. These data clearly oppose our hypothesis, that restoring the lack of PPE-MPTR immune responses in BCG increases its protective efficacy.

**Figure 6:**
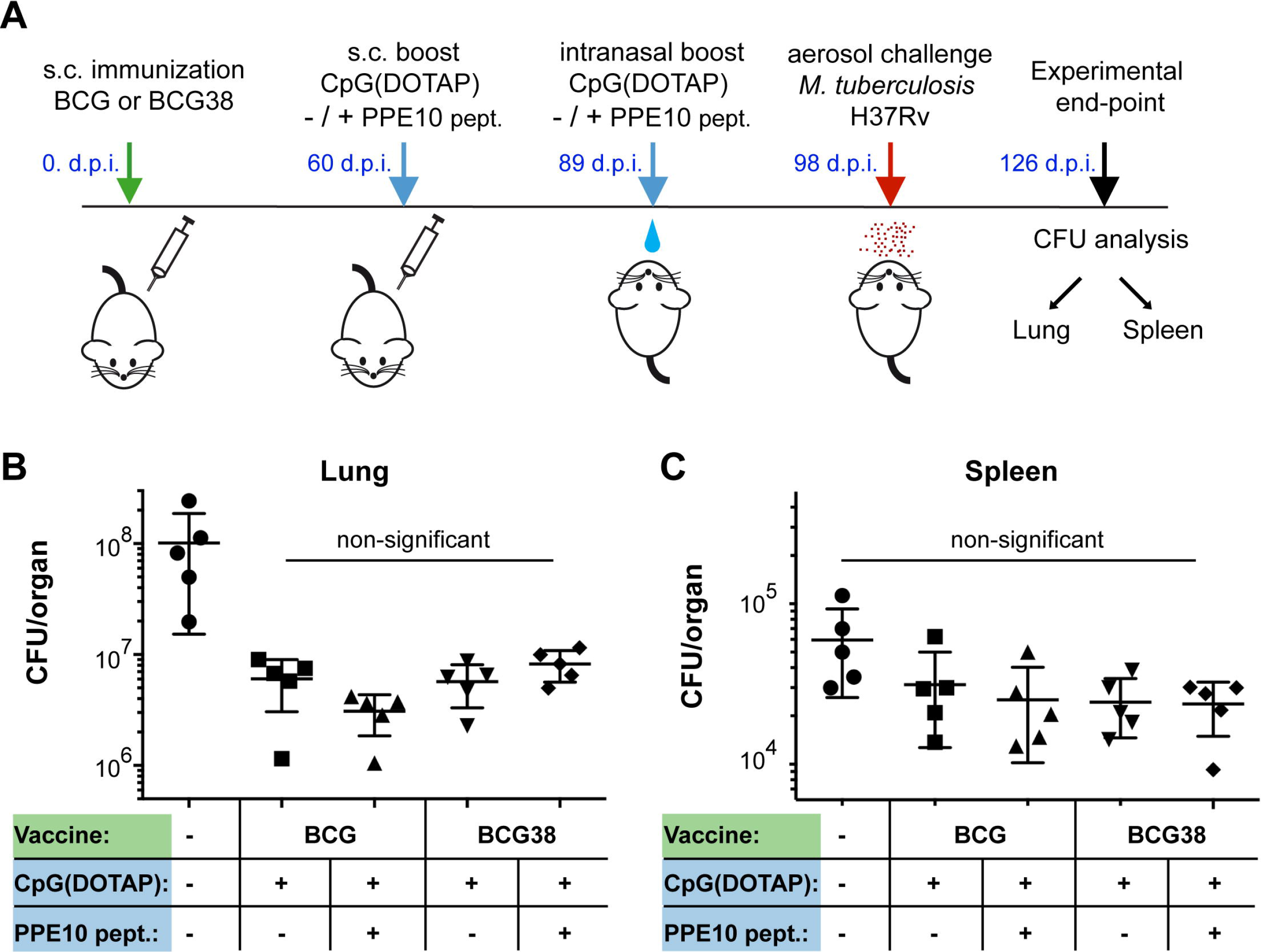
Boosting PPE10 specific immune responses does not increase protection against *M. tuberculosis*. A) Graphical representation of the prime-boost vaccination protocol. Mice were immunized with either BCG or BCG38 (Green). 60 days post-infection (d.p.i.) C57BL/6 x CBA F1 mice were injected s.c. with a booster consisting of adjuvant CpG(DOTAP), alone or in combination with a mix of PPE10_221-235_ and PPE10_381-395_ peptides (blue). The same formulation was intranasally administered four weeks later. Nine days after the intranasal boost, mice were exposed to *M. tuberculosis* H37Rv aerosol infection (220 CFU/lung 1 d.p.i). Bacterial lung (B) and Spleen (C) burdens were assessed by dilution and counting 4 weeks post-infection (experimental end-point) after being photographed for macroscopic investigation (Supplemental figures 2C, D). Each data point represents the CFU value of one organ from a single mouse, error bars depict the standard deviation. No significant differences between the vaccination conditions were detected by ordinary one-way ANOVA followed by Tukey’s test of multiple comparisons. All vaccination conditions resulted in a significant (*p*<0.01) reduction in lung burden compared to unimmunized control. Reduction in spleen CFUs was not significant for any of the vaccination conditions. Statistical analyses were performed using PRISM software.

Together, we could find no evidence of an immunomodulatory effect of PPE38-dependent proteins. Inversely, restoration of BCG’s capacity to secrete PE_PGRS and PPE-MPTR proteins and thereby enlarging the PE_PGRS/PPE-MPTR antigenic repertoire of BCG, did not result in improved vaccine protection in two mouse models.

## Discussion

We previously demonstrated that loss-of-function mutations in the *ppe38-locus* of *M. tuberculosis* block PE_PGRS and PPE-MPTR secretion and increase virulence in a mouse model [37]. In this work, we examined the correlation of known *ppe38* deletions in other lineages of the MTBC with a PE_PGRS/PPE-MPTR secretion defect. We hypothesized that the success of certain clinical isolates of Lineage 4, could perhaps be explained by their RD5-like deletion, which includes *ppe38* [46,77]. However, secretion analysis of these Lineage 4 strains revealed that a single copy of PPE71 carrying a the MGGAGAG-deletion, is still functional and sufficient to support PE_PGRS secretion. Similarly, although intriguing differences in protein secretion levels were observed between *M. canettii* strains, we found that all analyzed strains secreted PE_PGRS proteins. The anticipated polymorphisms in the ppe38-locus of selected *M. canettii* strains [43] were likely caused by a sequence assembly problem of repetitive sequences. These results highlight the difficulties of bio-informatic analyses of this locus, which is hampered by the high sequence similarity between *ppe38* and *ppe71*, that seem to cause already some discrepancies between the reference genomes of *M. tuberculosis* H37Rv and CDC1551 [29,37,38,78].

In contrast, our investigation of RD5-like polymorphisms did reveal that multiple members of the animal adapted lineage of the MTBC are completely devoid of PE_PGRS secretion because of their RD5 deletion. It should be emphasized that the RD5-like deletion of *M. orygis* occurred independently of that of *M. bovis* and *M. caprae*. Furthermore, even more members of the animal adapted lineage, such as *M. microti, M. suricattae* and the Dassie Bacillus, are reported to have independent RD5-like deletions, which we hypothesize to also block PE_PGRS and PPE-MPTR secretion [38,79–81]. Together, these findings suggest a specific selective advantage associated to loss of the *ppe38*-locus and its associated secretion phenotype in certain animal adapted strains. The modern Beijing strains, also defective in PPE38-dependent secretion, have expanded concurrently with increased human population densities and mobility [82]. These changes in the host-population alter the optimal balance between virulence/infectivity and lower the advantage to stay dormant or subclinical in the host [83]. It is tempting to speculate that the loss of PPE38 and its associated secretion and virulence phenotype has helped ancestral *M. tuberculosis* strains derived from human hosts, to adapt towards survival and transmission in a new host niches.

We were surprised that we were able to restore the secretion defect of BCG by introducing the ppe38-locus from *M. tuberculosis*. Since the RD5 deletion of BCG already occurred in the most-recent common ancestor of *M. bovis* and *M. caprae*, this deletion likely dates back millennia [47]. Furthermore, the 13 years of *in vitro* culturing by Calmette and Guérin to create BCG and the ensuing decades of culturing while it was distributed worldwide has caused accumulation of even more mutations [6,10,11]. Still, introduction of the integrative vector constitutively expressing the *ppe38-locus* was clearly able to restore both PE_PGRS and PPE-MPTR secretion in BCG.

Our newly identified immunogenic epitopes in the PPE-MPTR protein PPE10, provide a tool to gain more understanding about this group of proteins. Firstly, although previous work only definitively detected the C-terminal domain of PPE10 to be secreted [25,32,37], immunization with *M. tuberculosis* also clearly induced immune responses against the MPTR-associated epitope. This provides evidence that the MPTR domain is accessible to the host antigen presentation machinery and that these repetitive domains have the potential to contain functional T-cell epitopes. Furthermore, wild-type BCG and *M. tuberculosis* with impaired PPE38-dependent secretion were completely unable to induce immune responses against PPE10, similar to a general ESX-5 secretion mutant. This is important evidence that PPE38 is essential for the translocation of PPE-MPTR proteins through the ESX-5 secretion machinery *in vivo* and that without PPE38, these proteins are not surface associated or otherwise accessible to the immune system.

It is perhaps striking that the PE_PGRS and PPE-MPTR secretion defect of BCG has not been previously reported, considering the amount of research done on this vaccine. Based on the available literature on PE_PGRS and PPE-MPTR proteins, it is logical to hypothesize that a vaccine strain that does not secrete these proteins might in fact be a relatively effective vaccine. Many immunomodulatory properties have been attributed to PE_PGRS and PPE-MPTR proteins [14,33,34,62]. Perhaps the most relevant of these, is the reported function of certain PE_PGRS proteins to inhibit antigen presentation [35,36]. If PE_PGRS proteins indeed inhibit antigen presentation, it would be highly detrimental to introduce a vaccine that secretes these proteins. Notably, this is an urgent question since a number of novel tuberculosis vaccine candidates based on attenuated *M. tuberculosis* are currently in clinical or pre-clinical development. We showed for one of these candidate vaccines (*i.e. M. tuberculosis*-Δ*ppe25-pe19*), that PE_PGRS and PPE-MPTR secretion is indeed fully functional [52,63,71]. Our isogenic Δ*ppe38-71* strains of *M. tuberculosis* and the BCG38 strain form an ideal tool to answer such questions and to understand more about these proteins as a group. In this work, we did not find any evidence of inhibition of antigen presentation in strains secreting PPE38-dependent substrates, or lack thereof in strains without PPE38. Similarly, and in contrast to many reports of immunomodulatory effects of PE_PGRS proteins, we did not find any evidence of differential immune modulation by strains with, or without, functional PPE38-dependent secretion. More specifically, no differences were observed in DC maturation [84], MHC-I or -II expression [56] or cytokine production [85–87]. Finally, PE_PGRS and PPE-MPTR proteins have often been implicated as mycobacterial virulence factors [14,34,35,88,89]. The previously described increased virulence in strains lacking PPE38-dependent secretion, including the hypervirulent Beijing isolates, put this work in perspective [37]. Here, we bolster our previously published evidence that strains without PPE38, including a number of animal adapted species and the BCG vaccine, are truly unable to translocate these proteins. Although many of these animal adapted strains have reduced virulence in humans compared to *M. tuberculosis*, they are clearly pathogenic for their natural host and should not be seen as attenuated [90]. This is in line with a role for PPE38-dependent substrates as virulence attenuating factors [37]. Therefore, the biological roles of the PE_PGRS and PPE-MPTR proteins that are reported to be required for virulence, may not require secretion of these effector proteins or might in certain cases be due to indirect effects on other proteins. This hypothesis is further supported by the fact that many of the studies that attribute virulence traits to PE_PGRS and PPE-MPTR proteins, are performed in *M. smegmatis*, which lacks an ESX-5 secretion system and is unable to secrete these proteins [27,33,72]. Further work on the biological function of PE_PGRS and PPE-MPTR proteins, either on an individual basis or grouped, will have to take into account these findings and critically assess the impact of localization on effector function.

Perhaps the most relevant finding of this work is that BCG is unable to secrete PE_PGRS and PPE-MPTR proteins and therefore does not raise T-cell responses against these proteins. Previous studies have shown that antibodies can be raised against PE_PGRS proteins, suggesting that it could be a beneficial property of a vaccine to secrete these proteins [25,91,92]. Here we provide evidence that PPE-MPTR proteins can be immunogenic in mice, which is further supported by a recent publication investigating immunogenicity of the PPE-MPTR protein PPE39 [67]. Kim et al. identified two immunogenic epitopes of which one (MTBK_24820_85-102_) is located in the PPE-domain and has high homology to non-MPTR PPE proteins, while the other (MTBK_24820_217-234_) was located in the MPTR domain of this protein. Interestingly, the authors reported that vaccination with the recombinant PPE39 protein induced a higher level of protection against *M. tuberculosis* Erdman, compared to a hypervirulent Beijing isolate [67]. This difference could be explained by our data, which would suggest immune responses against that MPTR epitope would not be helpful against a PPE38-deficient Beijing isolate. A related issue that requires further work is whether the PPE38-dependent secretion effect in modern Beijing isolates is somehow related to that of the BCG vaccine and whether their respective secretion defects affect vaccine efficacy.

There is strong evidence for the importance of PPE-MPTR proteins in human immune responses, because the PPE-MPTR protein PPE42 (Rv2608) is an integral part of the subunit fusion-protein vaccine candidate ID93 [60,66]. The fusion protein ID93 consists of four different proteins and has been tested as a vaccine candidate in both a Phase 1 and Phase 2A clinical trial [93,94]. Bertholet et al. 2008 demonstrated that PBMCs isolated from PPD_+_ healthy subjects produced IFN-γ in response to PPE42 and that almost 70% of subjects showed a reaction against the recombinant protein in a recall experiment [66]. Interestingly, 100% of PPD_+_ subjects exhibited recall responses against the other (non-MPTR) PPE proteins that were tested, which could possibly be explained by exposure to modern Beijing, or other PPE38-deficient strains, in the subject cohort. PPE42 was selected as part of the ID93 vaccine due to its excellent ability to induce both humoral and cellular immune responses and immunization with PPE42 provided protection in mice almost comparable to BCG [60,66]. In Guinea pigs, ID93 significantly boosted the protection induced by BCG, which was interpreted as an ability to boost immune responses elicited by BCG [60]. However, based on our work it should be assumed that BCG does not induce immune responses against the PPE-MPTR protein PPE42 and that boosting with ID93 may in fact broaden antigenic repertoire of the combined vaccination. Similarly, ID93 is able to induce protective immune responses to the *M. tuberculosis* Beijing isolate HN878, but it is unclear what the role of PPE42 is in this response. The analyses performed in Bertholet et al. 2010 and Baldwin et al. 2015 were performed with the four-gene fusion protein ID93 and not with the individual PPE42 subunit, which makes it impossible to assess these questions more thoroughly. What remains clear however, is that the PPE-MPTR protein PPE42 is an important part of a vaccine currently in clinical trials. The finding that ID93 includes a protein to which parental BCG is likely not able to induce immune responses, may actually put the proven booster qualities of this vaccine candidate in a different light and lead to optimal strategies to employ it.

The question whether immune responses against PPE38-dependent proteins are important for a vaccine to be protective against tuberculosis, needs an urgent answer, especially since it concerns a total of 89 proteins. There are multiple vaccine candidates in clinical, or pre-clinical, development that are based on attenuated *M. tuberculosis* strains and which likely secrete PE_PGRS and PPE-MPTR proteins [24,52,62,63,71,95]. Should we knock-out *ppe38-71* in these vaccine candidates to avoid immune modulation by the secreted substrates, or should we prioritize these vaccine candidates, because they have a broader potential repertoire of epitopes? Should BCG vaccination be boosted by vaccine candidates including PPE-MPTR proteins such as ID93, or should this be avoided? Are there differences between designing vaccine candidates against strains secreting PE_PGRS/PPE-MPTR proteins and those with a PPE38-dependent secretion defect, such as the modern Beijing isolates? Are murine or other small animal infection models appropriate to predict PE_PGRS and PPE-MPTR-mediated impact on vaccine efficacy? These are questions that we are not yet able to answer in this work, but they reveal the need to increase our understanding of PE_PGRS and PPE-MPTR proteins. Better knowledge on PE_PGRS/PPE-MPTR proteins is not just an intellectual goal, but may also help to make more informed decisions in the design of novel vaccines against tuberculosis.

## Acknowledgements

The authors would like to thank Alexandre Pawlik and Fiona McIntosh for help. We also thank Michal Brennan for initially sharing a clone producing PE_PGRS antibody. We thank Robyn Lee and Anzaan Dippenaar for data analysis. We thank James Gallant, Maroeska Burggraaf and Edith NG Houben for insightful discussions.

## Ethical approval

Studies in immunocompetent mice were performed according to European and French guidelines (Directive 86/609/CEE and Decree 87–848 of 19 October 1987) after approval by the Institut Pasteur Safety, Animal Care and Use Committee (Protocol 11.245) and local ethical committees (CETEA 2012–0005 and CETEA 2013–0036).

## Figure Legends

**Supplemental Figure 1: Immunoblot secretion analysis reveals no PPE38-dependent secretion effect in *M. canetti*, or *M. tuberculosis* Mj-sublineage Lineage 4 strains affecting the Nunavik Inuit.** Immunoblots of whole-cell lysates or culture filtrates of the indicated *M. canetti* (A), *M. tuberculosis* (B) or BCG (C) isolates [43,46]. A) Although differences in protein secretion could be observed between different *M. canetti* isolates (A-J), all isolates exhibited PE_PGRS secretion. B) PE_PGRS secretion of Mj-sublineage strains with a deletion affecting *ppe38*, but not *ppe71* (Lanes 4-8) was not discernible from Lineage 4 control isolate CDC1551 or an isolate from the same cohort without this deletion (MT13848). C) Introduction of plasmid pMV::*ppe38-71* in BCG complemented PE_PGRS secretion (BCG38), while complementation was not observed when performed with pYUB::RD5, even though presence of genetic presence of RD5 was PCR-confirmed with primers RD5B-plcA.int.F/R [47]. Anti-SigA staining is uses as a lysis control in A, while anti-GroEL2 is used in B and C. Strain details can be found in Supplemental Table 5. Full blots of panels A-C are depicted in Supplemental Figure 6.

**Supplemental Figure 2: Harvested lungs and spleens from vaccinated and *M. tuberculosis* infected mice.** Organs depicted in A and B correspond to the experiment depicted in Figure 3. Organs depicted in C and D correspond to the experiment depicted in Figure 6. After photography of the lungs (A, C), a single lung lobe was used for lung CFU quantification. Splenomegaly (B, D) was reduced, by all vaccination conditions, but did not differ markedly between vaccination conditions.

**Supplemental Figure 3: Epitope mapping of a peptide library identifies two PPE10 epitopes that are immunogenic in C57BL/6 x CBA mice.** IFN-γ production in response to peptides covering the indicated amino acid positions of PPE10 (Rv0442c) in C57BL/6 (grey/green) or C57BL/6 x CBA F1 (B6CBAF1, blue/brown) mice. Mice were immunized with *M. tuberculosis* H37Rv (left) or unimmunized (right).

**Supplemental Figure 4: Construction and secretion analysis of *M. tuberculosis* CDC1551-Δ*ppe10*.** A) Schematic representation of deletion strategy and primers. The genetic region around PPE10, as taken from tuberculist, is depicted in colored arrows [69]. Flanking fragments used for homologous recombination are depicted in black bars. Left (PPE10KO-LF and PPE10KO-LR) and right (PPE10KO-RF and PPE10KO-RR) flanking regions were amplified by primers depicted in black. Primers used to verify successful homologous recombination are depicted in dark blue. All primer sequences can be found in Supplemental Table 4. B) PCR verification of successful homologous recombination in seven different colonies that grew on hygromycin selection plates. Colony 1 was taken for further analyses. C) Immunoblot analysis of strains tested for immunogenicity in Figure 5. Samples were prepared as described in materials and methods section. SigA was used as a loading in lysis control. Some lysis could be found in both BCG and BCG38, but was not markedly different between strains. Full gels and blots used to create B and C are depicted in Supplemental Figure 7

**Supplemental Figure 5: Full blots corresponding to panels depicted in Figure 1B-C.**

**Supplemental Figure 6: Full blots corresponding to panels depicted in Supplemental Figure 1A-C.**

**Supplemental Figure 7: Full blots and gels corresponding to panels depicted in Supplemental Figure 4B-C.**

**Supplemental Table 1:** Mean Fluorescent Intensities and percentage of live C57BL/6 BM-DCs, infected (MOI = 0.5) with indicated strains of *M. tuberculosis* or *M. bovis* BCG with or without the *ppe38*-locus. As expected, the percentage of live cells was higher for BCG-infected cells than for cells infected with *M. tuberculosis* strains, but did not vary significantly between isogenic strains (≤ 1.0 % difference between isogenic strain). These values are derived from the experiment depicted in Figure 2A.

|  | CD40 | CD80 | CD86 | H-2K<sup>b</sup> | I-A<sup>b</sup> | % Live<br>Cells |
| --- | --- | --- | --- | --- | --- | --- |
| **Uninfected** | 218 | 338 | 264 | 663 | 904 | 94.9 |
| **CDC1551** | 282 | 449 | 427 | 982 | 1404 | 82.5 |
| **$\Delta ppe38-71$** | 287 | 424 | 446 | 925 | 1360 | 81.7 |
| ***ppe38-71-C*** | 303 | 457 | 454 | 969 | 1419 | 81.5 |
| **BCG** | 273 | 430 | 390 | 810 | 1348 | 86.6 |
| **BCG38** | 276 | 429 | 381 | 832 | 1417 | 86.8 |

**Supplemental Table 2:**
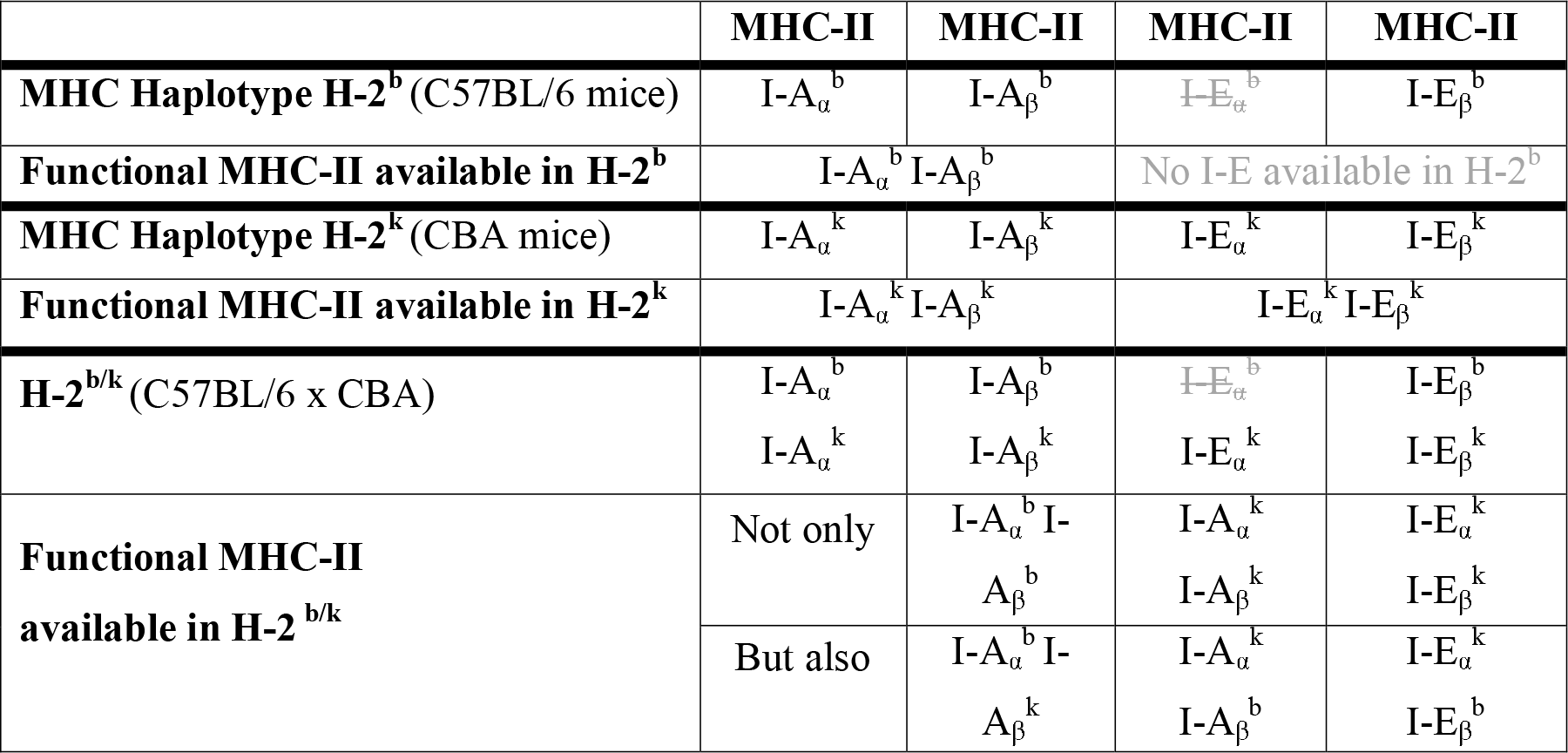
Why there are more MHC-II restricting molecules available in C57BL/6 x CBA F1 mice than in C57BL/6 or CBA mice. A promoter mutation disrupts production of I-Eα^b^ in C57BL/6 mice (Grey font with strikethrough), which are therefore unable to produce MHC-II I-E (Grey). In contrast, H-2^k^ mice can produce both I-Aα^k^ I-Aα^k^ and I-Eα^k^ I-Eα^k^ C57BL/6 x CBA F1 mice have an even bigger repertoire of possible functional MHC-II isoforms available due to recombination between the subunits.

**Supplemental Table 3:**
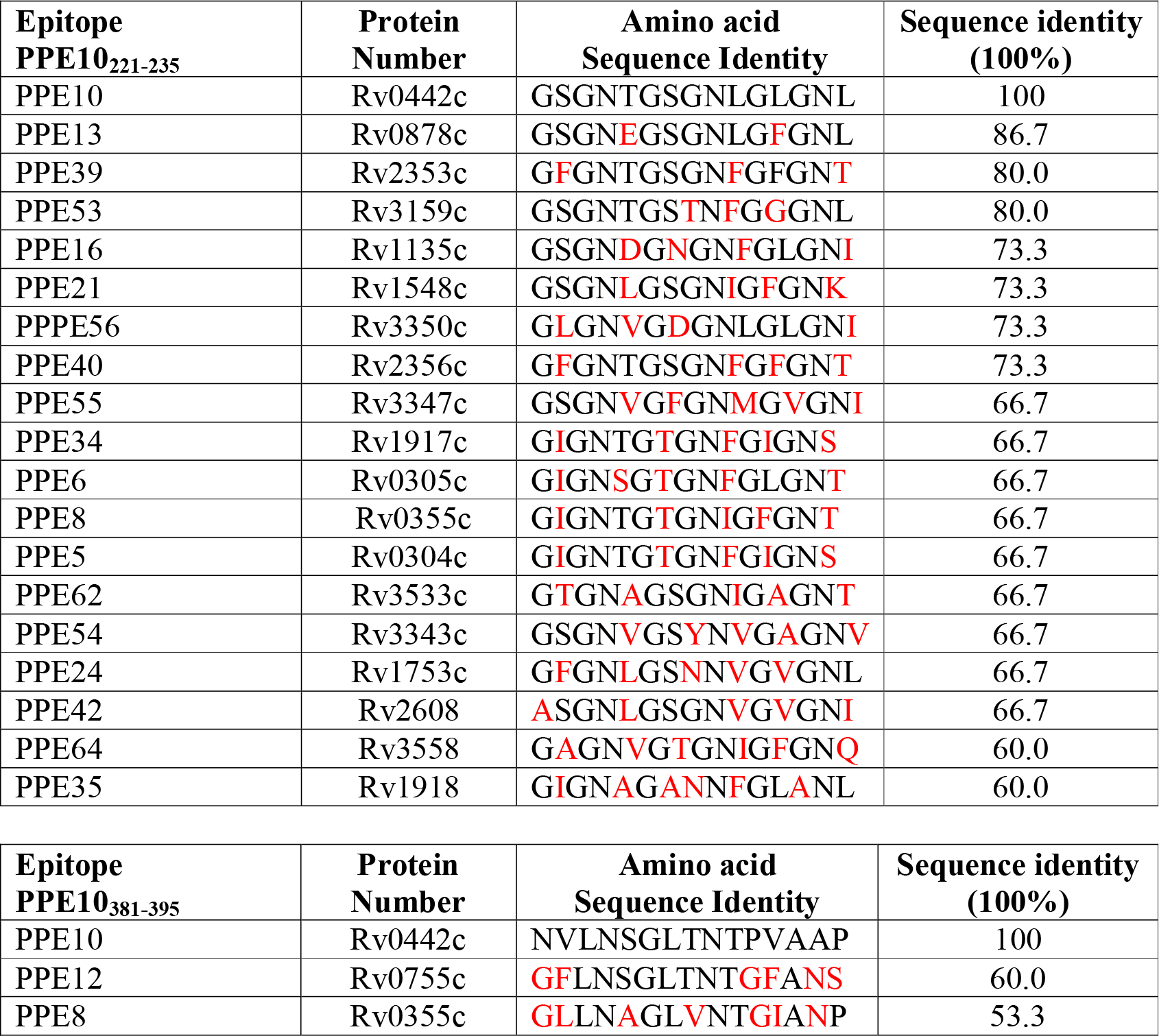
Sequence identity determined by BlastP search of immunogenic epitopes against the genome of *M. tuberculosis* H37Rv [29,96]. Black letters indicate identical amino acids. Red letters indicate non-identical amino acids. Top: homologues of the MPTR-containing peptide PPE10_221-235_ ordered by percentage of sequence identity. Bottom: Homologues of the peptide PPE10_381-395_, which is part of the C-terminal secreted domain of PPE10.

**Supplemental Table 4:**
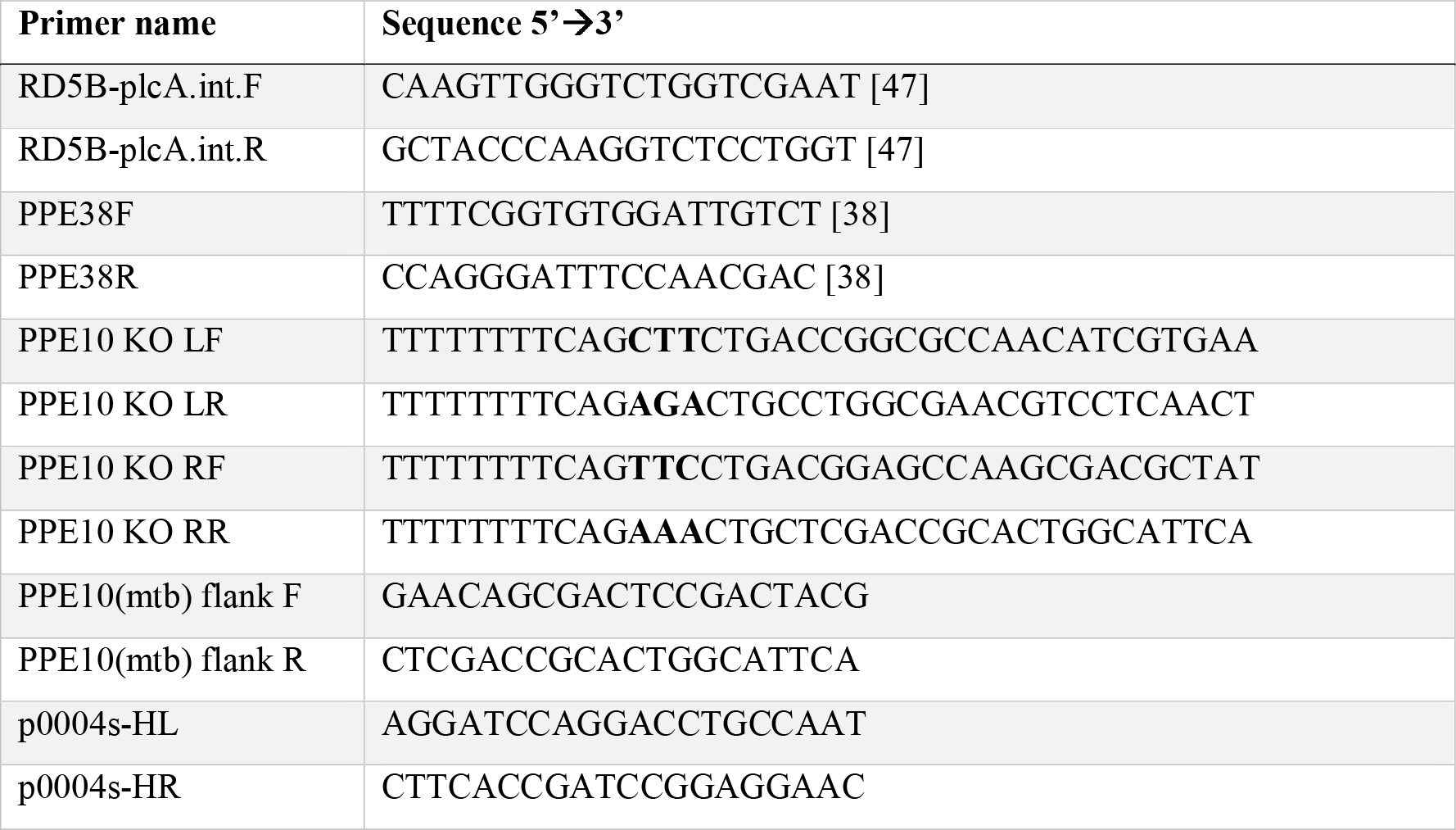
Primers used in this study

**Supplemental Table 5:**
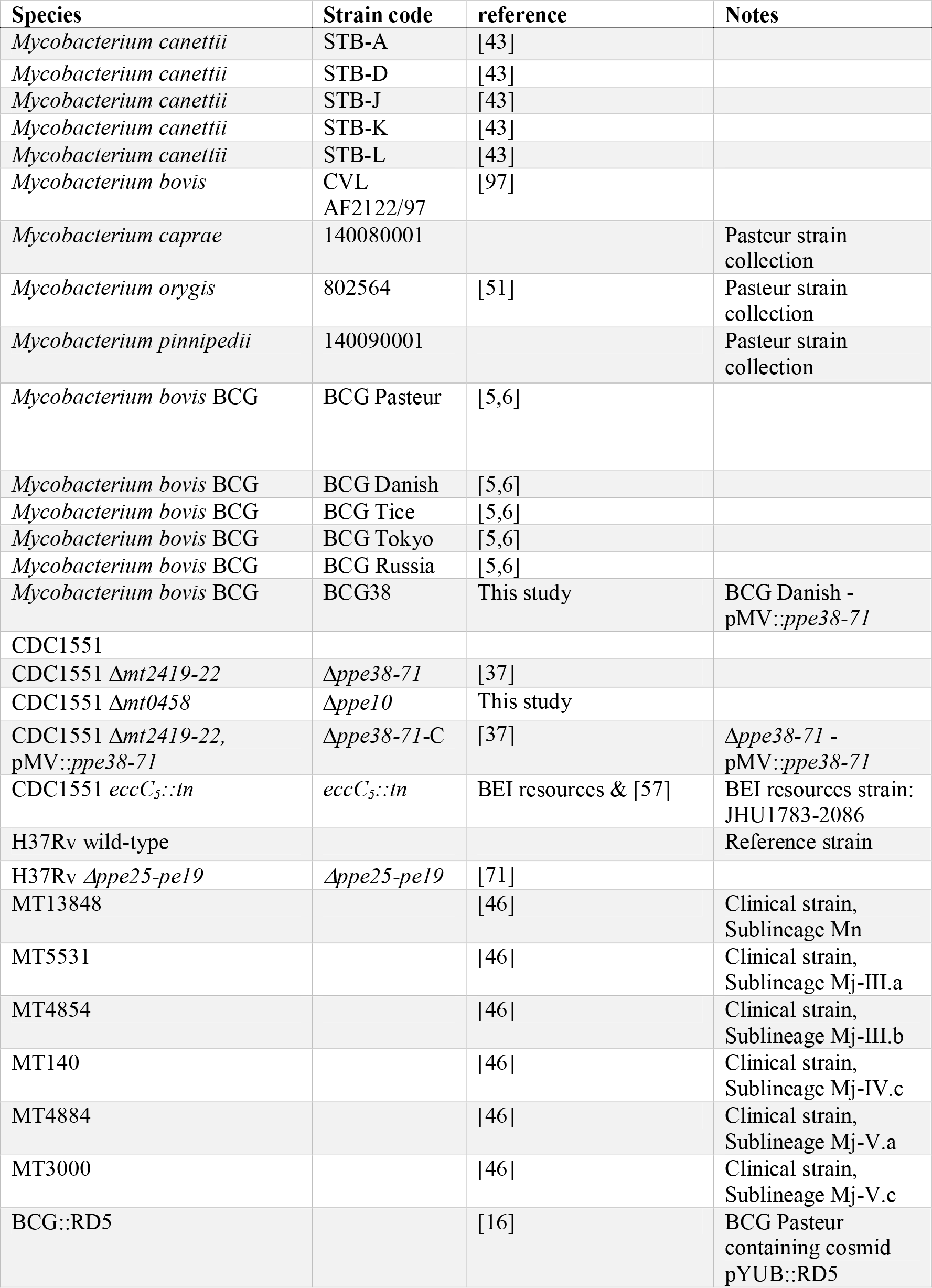
Bacterial strains used in this study

## Materials and methods

### Strains and growth conditions

All strains used in the study and the sources they are derived from can be found in supplemental table 5. Unless otherwise specified, all mycobacterial strains were grown on Middlebrook 7H11 solid medium (Difco) supplemented with OADC (BD Biosciences), or liquid 7H9 medium supplemented with ADC supplement and 0.05% Tween-80. Antibiotics were added where opportune at a concentration of 50µg/ml for Hygromycin (Euromedex), or 25µg/ml for Kanamycin (Sigma). Strains were incubated at 37°C. Liquid cultures were grown in shaking conditions at 80 rotations per minute. For animal-adapted strains *M. bovis, M. caprae, M. orygis* and *M. pinnipedii*, 0.2% w/v of Pyruvate (Sigma) was added to the growth medium [98]. Infection stocks of *M. tuberculosis* H37Rv used for aerosol infection experiments and BCG or BCG38 vaccination stocks without Tween-80 were prepared by inoculating 0.1 OD/ml bacteria in 100ml liquid culture without Tween-80. This culture was incubated for 7 days, after which it was washed with phosphate buffered saline (PBS) and sonicated (5x (100 pulses of 0.1s)) and left to rest for at least one hour before collecting the cell suspension considered to obtain a single-cell solution of encapsulated mycobacteria. Standard vaccination stocks were prepared in Dubos medium containing 0.025% Tween-80 in standing conditions and were harvested at an optical density between 0.4 and 0.7 OD_600_/ml.

### PCR verification of RD5 deletions

RD5 deletions were PCR verified by previously published primers specific for *plcA* (*rv2351c* - Supplemental Table 4), which produce a product of approximately 500bp when this gene is present [47]. Primers amplifying the *ppe38-71*-locus (Supplemental Table 4) produce a 3378bp product when the complete *ppe38-71* locus is present [38]. This includes two copies of *ppe38/71* (*mt2419/mt2422*) flanking the *esxX* (*mt2420*) and *esxY* (*mt2421*) in between in CDC1551. When only one copy of *ppe38* and no *esxX/esxY* are present this PCR produces a product of approximately 1500 bp [38].

### Recombinant strains and mutant construction

The complementation plasmid containing the ppe38-locus from CDC1551 (*mt2419-22*) under expression of *hsp60* promoter was previously described [37]. The cosmid containing the RD5 region (pYUB::RD5) was part of the library described by Bange et al. 1999 and contains the genetic region spanning 2,611 kb – 2,645 kb of the *M. tuberculosis* H37RV reference genome [29,99].

*M. tuberculosis*-Δ*ppe10* was constructed as described by Bardarov et al. [70]. The homologous recombination construct was created by a PCR combining primers PPE10 KO LF & LR to amplify the 3’ end of *rv0442c* and another PCR with primers PPE10 KO RF & PPE10 KO RR to amplify the 5’ end of *rv0442c* (See Supplemental Table 4 for primer sequences). After phage packaging and infection, seven transformed colonies were tested by PCR with either primer PPE10(mtb) flank F & p0004s-HR, or PPE10(mtb) flank R & p0004s-HL (Supplemental Figure 4A, B). All colonies were found to have the correct deletion spanning from 152bp to 1133bp after the 5’ of *rv0442c*. We attempted to complement the Δ*ppe10* mutant with a previously published plasmid (p19kPro::*rv0442c*-HA) overexpressing HA-tagged PPE10 under control of the *1pqH* promotor [25]. Although clones expressing the HA-tag on this plasmid were obtained, these had a considerable *in vitro* growth defect, which would conflict with *in vivo* and *in vitro* studies and therefore this complemented strain was not analyzed further.

### Secretion analysis

Strains were pre-cultured until mid-logarithmic phase under normal growth conditions (described above). Cultures were washed two times in 7H9 medium without ADC, supplemented with 0.2% Dextrose and 0.05% Tween and were incubated in this medium for 48 hours. Cultures were centrifuged to separate cells and the supernatant was filtered through a 0.02µm filter, after which it was TCA-precipitated to concentrate. Cellular material was washed with PBS, resuspended in solubilisation/denaturation buffer and boiled for 10 min at 95°C. After sterilisation by heating for 2 hours at 80°C, samples were sonicated to disrupt cells and boiled at 95°C during 10 minutes.

Samples were loaded on 12% or 4-12% SDS-Page gels (NuPage ®, Novex, Life technologies) and transferred to nitrocellulose filters by dry western blotting (iBlot ®, Invitrogen). Proteins were stained by primary antibodies: Anti-PGRS 7C4.1F7 [25] (Clone 7C4.1F7 was a kind gift from Michael J. Brennan, USA), polyclonal anti-SigA (Kind gift from I. Rosenkrantz, Denmark), Rabbit polyclonal anti-EsxN (rMTb9.9A) [100], monoclonal ESAT-6 (hyb76-8), or anti PPE41 [101].

### Cell infection, ELISA and flow cytometry

BM-DCs derived from C57BL/6 (H-2^b^) female mice were generated directly in 6-well plates and infected at day 6 of culture with different mycobacterial strains at M.O.I of 0.5 in RPMI 1640-GlutaMax medium (Invitrogen) containing 10% FBS (4 × 10^6^ cells/well in 4 ml volume). After over-night of infection at 37°C and 5% CO_2_, IL-6 (clone MP5-20F3 for coating and clone MP5-32C11 for detection, BD Pharmingen), IL12p40/70 (clone C17.8 RUO, BD Pharmingen) and TNF-α (clone 1F3F3D4 for coating and clone clone XT3/XT22 for detection, eBioscience) cytokine production was quantified in the culture supernatants by ELISA.

For viability and phenotypic maturation evaluation, infected DCs were washed with PBS and incubated first with Live/Dead-Pacific Blue reagent (Invitrogen) during 35 minutes at 10°C in the dark. Cells were then washed twice and incubated with appropriate dilution of anti-CD16/CD32 (2.4G2 mAb, BD Pharmingen) during 20 minutes followed by surface staining by 30 minutes of incubation with appropriate dilutions of APC-anti-CD11b (BD Pharmingen), PE-Cy7-anti-CD11c (BD Pharmingen), FITC-anti-CD40 (clone HM40-3, SONY), FITC-anti-CD80 (B7-1) (clone 16-10A1 Biolegend), FITC-anti-CD86 (B7-2) (clone PO3, SONY), FITC-anti-MHC-II (I-A/I-E) (clone MS/114.15.2, eBioscience), FITC-anti-MHC-I (H-2k^b^) (clone AF6-88-5-5-3, eBioscience) or FITC-anti-IgG1k isotype control. The stained cells were washed twice with FACS buffer (PBS containing 3% fetal bovine serum (FBS) and 0.1% NaN_3_) and then fixed with 4% paraformaldehyde during 18h at 10°C prior to sample acquisition by a LSR Fortessa flow cytometer system (BD Bioscience) and BD FACSDiva software. The obtained data were analyzed using FlowJo software (Treestar, OR, USA).

### Antigen presentation assay

BM-DCs derived from BALB/c (H-2^d^) female mice were used at day 6 of culture as antigen presenting cells. Cells were seeded in 96-well plates at 5 × 10^4^ cells/well and loaded with 1 µg/ml of homologous or negative control synthetic peptides, or infected with different mycobacterial strains with serial two-fold dilutions of M.O.I., starting at M.O.I. = 10, in RPMI 1640-GlutaMax medium (Invitrogen) containing 10% FBS. After 18h of infection at 37°C and 5% CO2, cells were washed twice with RPMI medium to eliminate the IL-2 possibly produced by the infected DCs and then co-cultured with 1 × 10^5^ cells/well of T-cell hybridoma specific to EsxH/TB10.4_74-88_ (1G1) or Ag85A_101-120_ (2A1), respectively restricted by I-A^d^ or I-E^d^. After over-night of co-culture at 37°C and 5% CO_2_, the IL-2 secretion was quantified in the culture supernatants by ELISA (clone JES6-1A12 for coating and clone JES6-5H4 for detection, BD Pharmingen).

### Epitope mapping of PPE10 and T-cell assay

A peptide library of sixty 15-mers with a 5-amino acid shifting frame, spanning amino acids 181-487 of PPE10 (Rv0442c), was constructed commercially (Mimotopes Europe, United Kingdom). Epitope screening of PPE10 and immunogenicity assays were performed as previously described [52], with some modifications. Briefly, 6-8-week-old female C57BL/6 (H-2^b^) or C57BL/6 x CBA F1 (H-2^b/k^) mice were immunized s.c. with 1 × 10^6^ CFU/mouse of different mycobacterial strains obtained from exponential culture in Dubos medium. Epitope mapping was performed with mice immunized with *M. tuberculosis* H37Rv. Three to four weeks post-immunization, mice were sacrificed and pool of total splenocytes (*n* = 2 mice per group) were restimulated in 96-well flat-bottom plates (TPP, Den- mark) at 5 × 10^5^ cells per well in HL-1 medium (Biowhittaker, Lonza, France), complemented with 2 mM GlutaMax (Invitrogen, Life Technologies, France), 5 × 10^5^ M β-mercaptoethanol, 100 U/ml penicillin and 100 μg/ml streptomycin (Sigma-Aldrich, France) in the presence of 10-20 μg/ml of individual peptides. IFN-γ production in the supernatant was quantified by ELISA after 72h of culture at 37°C and 5% CO2 (clone AN-18 for coating and clone R46A2 for detection), BD Pharmingen.

### Protection assays

BCG and BCG38 were grown in 10ml Dubos medium or in 100ml 7H9-medium with ADC-supplement without Tween-80. *M. tuberculosis* H37Rv and BCG-strains cultured without Tween-80 were sonicated (5 × 100 pulses; 0.1 seconds/pulse; 0.9 seconds’ rest; amplitude 30%) to disrupt clumps and were frozen at −80°C. Frozen stocks were counted for CFU’s before immunization to assess dose while the dose of Dubos-grown strains was estimated based on optical density.

Eight-week-old C57BL/6 mice (n = 5 mice/group), were immunized with 1 × 10^6^ CFU/mouse of BCG Danish (cultured − or + Tween-80), or BCG38 (cultured − or + Tween-80) in 200 μl PBS. Eight mice were concurrently injected with sterile PBS. Thirty days after vaccination, mice were challenged with aerosolized WT *M. tuberculosis* H37Rv strain. Three mice were sacrificed to assess bacterial lung burdens 1 day post challenge (assessed at 680 CFU/lung). All other mice were killed four weeks post-challenge due to human end-point criteria of unvaccinated mice. Lungs and spleens were homogenized by beadbeating, serially diluted in PBS and plated on 7H11 plates with (Lungs) or without (spleens) BBL™ MGIT™ PANTA™ (Beckton Dickinson, Ireland).

The Prime-boost vaccination and challenge experiment was performed similar as above, with the following modifications. BCG or BCG38 were precultured in Dubos medium and five first generation C57BL/6 x CBA crossover mice were left unvaccinated or s.c. immunized (*n* = 5 mice/group). Eight weeks post-immunization, a subcutaneous boost was administered. This boost consisted of 200 μl/mouse of formulation containing 50 μl of each PPE10-derived peptide (PPE10_221-235_ and PPE10_381-395_) ProteoGenix, France, 30 μg of CpG 1826 oligodeoxynucleotides as adjuvant (Sigma-Aldrich, France) at 1 μl/mL concentration, 60 μl of liposomal transfection reagent DOTAP (N-[1-(2,3-DioleOyloxy)]-N,N,N-Trimethyl Ammonium Propane methylsulfate, Roche, France) and 10 μl Opti-MEM (Life Technologies, France) as described in Sayes et al., 2016 [63]). Four weeks later, an intranasal boost was given to mice via intra-nasal route, under anesthesia as described in Sayes et al., 2016, 25 μl/mouse contained 10 μg of PPE10 peptides, 2 μg of CpG at 10 μl/mL concentration, 10 μl of DOTAP and 3 μl Opti-MEM contained in 20 μl/mouse [63]. Ten days after the intranasal boost, mice were aerosol challenged with WT *M. tuberculosis* H37Rv strain. Three non-immunized mice were killed one day post challenge to assess infectious dose administered, which was calculated at 220 CFU/lung. Four weeks later all other mice were killed and one lung and the spleen were homogenized with a MillMixer organ homogenizer (Qiagen, Courtaboeuf, France) and plated to assess bacterial burdens on 7H11 Agar medium supplemented with ADC (Difco, Becton Dickinson). The CFU were counted after 3-4 weeks of incubation at 37°C.

All immunized and infected mice for immunogenicity and protection experiments were placed and manipulated in isolator in BSL-III protection-level animal facilities at the Pasteur Institute.

To determine the statistical significance of the data, analyses were performed by use of GraphPad Prism software (GraphPad Soft-ware, La Jolla, CA, USA), using ordinary one-way ANOVA followed by Tukey’s test for multiple comparisons.

